# Heat Stress Interferes with Formation of Double-Strand Breaks and Homolog Synapsis in *Arabidopsis thaliana*

**DOI:** 10.1101/2020.10.02.324269

**Authors:** Yingjie Ning, Qingpei Liu, Chong Wang, Erdai Qin, Zhihua Wu, Minghui Wang, Ke Yang, Ibrahim Eid Elesawi, Chunli Chen, Hong Liu, Rui Qin, Bing Liu

## Abstract

Meiotic recombination (MR) drives novel combination of alleles and contributes to genomic diversity in eukaryotes. In this study, we showed that heat stress (36-38°C) over fertile threshold fully abolished crossover (CO) formation in Arabidopsis. Cytological and genetic studies in wild-type plants, and the *syn1* and *rad51* mutants suggested that heat stress reduces generation of SPO11-dependent double-strand breaks (DSBs). In support, the abundance of recombinase DMC1, which is required for MR-specific DSB repair, was significantly reduced under heat stress. In addition, we showed that high temperatures induced disassembly and/or instability of ASY4-but not SYN1-mediated chromosome axis. At the same time, ASY1-associated lateral element of synaptonemal complex (SC) was partially affected, while the ZYP1-dependent central element of SC was disrupted, indicating that heat stress impairs SC formation. Moreover, quantitative RT-PCR revealed that genes involved in DSB formation; e.g. *SPO11-1*, *PRD1*, *2* and *3*, were not impacted; however, recombinase *RAD51* and chromosome axis factors *ASY3* and *ASY4* were significantly downregulated under heat stress. Taken together, these findings revealed that heat stress inhibits MR via compromised DSB formation and homolog synapsis, which are possible downstream effects of the impacted chromosome axis. Our study thus provides evidence shedding light on how increase of environmental temperature influences MR in Arabidopsis.

## Introduction

Male meiosis is the basis for sexual reproduction and is important for genomic diversity and ploidy stability of plants over generations. Meiotic recombination (MR) allows reciprocal exchange of genetic information and assures balanced chromosome segregation at later meiotic stages (Wang and Copenhaver, 2018). MR is initiated by programmed formation of double-strand breaks (DSBs) catalyzed by SPO11, a conserved type II topoisomerase (topoisomerase VI, subunit A) among eukaryotes (Bergerat et al., 1997; Grelon et al., 2001). In Arabidopsis, there are three SPO11 homologues which share 20%-30% sequence identity (Hartung and Puchta, 2000, 2001). SPO11-1 and SPO11-2 are required for MR, while SPO11-3 plays a role in DNA endo-reduplication (Hartung et al., 2007; Stacey et al., 2006; Sugimoto-Shirasu et al., 2002). DSB formation is crucial for synapsis of homologous chromosomes and CO formation, with both the *spo11-1* and *spo11-2* mutants having omitted MR and impaired male fertility due to disordered chromosome segregation (Grelon et al., 2001; Stacey et al., 2006). Thereafter MRE11-RAD50-NBS1/XRS2 (MRN/X) complex generates 3’ single-strand DNAs (ssDNAs) at DSB sites, and recombinases RAD51 and DMC1 bind to the ssDNA overhangs to promote homologous searching and pairing (Hartung et al., 2007; Milman et al., 2009; Neale et al., 2005; Uanschou et al., 2007). DMC1 and RAD51 localize and function separately at DSB sites, with RAD51 acting as an accessory factor for DMC1 in catalyzing MR (Cloud et al., 2012; Da Ines et al., 2013; Kurzbauer et al., 2012; Lan et al., 2020). It was evidenced that RAD51 plays a role in non-crossover (NCO) pathway in DSB repair using sister chromatid as template, while DMC1 catalyzes meiotic-specific DSB repair (Cloud et al., 2012; Singh et al., 2017).

Chromosome axis and synaptonemal complex (SC) are synchronously built as initiation of DSB formation. In Arabidopsis, sister chromatids bound by meiosis-specific ɑ-kleisin AtRec8/SYN1 form a loop structure, the basal regions of which are connected to axial region of SC; and it is proposed that DSB formation is proposed to take place at the chromatin loops tethered to the axis (Lambing et al., 2020b; Zickler and Kleckner, 1999). Two coiled-coil proteins ASY3 and ASY4 are essentially required for axis formation (Armstrong et al., 2002; Chambon et al., 2018; Ferdous et al., 2012; Sanchez-Moran et al., 2008). Normal DSB formation and MR rely on construction of chromosome axis. Null mutation in AtRec8/SYN1, or disorganization of ASY3 and/or ASY4-mediated chromosome axis leads to reduced DSB formation and disrupted MR (Bai et al., 1999; Chambon et al., 2018; Ferdous et al., 2012; Lambing et al., 2020b). HORMA domain protein ASY1 and transverse filament protein ZYP1 mediate the formation of lateral and central element of SC, respectively, which are also required for synapsis of homologous chromosomes and CO formation (Armstrong et al., 2002; Barakate et al., 2014; Higgins et al., 2005; Wang et al., 2010). Interactions between ASY1 and ASY4, and ASY4 and ASY3 aid chromatin localizing to the lateral element of SC (Chambon et al., 2018; Yang et al., 2019). Moreover, ASY1 mediates the occurrence of DMC1-dependent MR, and it was recently shown to be essential for maintenance of CO interference (Lambing et al., 2020a; Sanchez-Moran et al., 2007).

Environmental temperatures can affect multiple processes of meiotic cell division in plants (Liu et al., 2019; Morgan et al., 2017). In Arabidopsis, wheat, Populus and persimmon, temperature stresses affect formation and organization of spindle and/or phragmoplast during male meiosis, which results in meiotic restitution or aneuploid gametes (De Storme et al., 2012; Lei et al., 2020; Liu et al., 2018; Mai et al., 2019; Tang et al., 2011; Wang et al., 2017). In the context of MR, both decrease and increase of temperature within fertility threshold (8-28°C) promotes MR rate via enhanced type-I CO formation in Arabidopsis (Francis et al., 2007; Lloyd et al., 2018; Modliszewski et al., 2018). A higher temperature at 32°C was found to reduce CO formation, probably by influencing synapsis of homologous chromosomes (De Storme and Geelen, 2020). However, how high temperatures beyond fertility-tolerable range influence MR, especially for early events; i.e. DSB and axis formation, remains unknown. In this study, we demonstrated that high temperatures beyond fertility threshold (36-38°C) fully inhibit CO formation in Arabidopsis. We showed that heat stress reduces SPO11-dependent DSB formation and affected the assembly and/or stability of ASY4- but not SYN1-mediated chromosome axis. We also showed that ASY1-mediated lateral element and ZYP1-dependent central element of SC were partially impacted, or disrupted, respectively. Overall, our findings revealed that heat stress interferes with MR via compromised DSB formation and homolog synapsis in *Arabidopsis thaliana*.

## Results

### High temperatures beyond fertility threshold fully inhibit CO formation

To reveal the impact of high temperature beyond fertility-tolerable range on MR, we analyzed chromosome spread in heat-stressed (36-38°C, 24 h) flowering wild-type Col-0 plants using 4′,6-diamidino-2-phenylindole (DAPI) staining. Under normal temperature conditions, leptotene chromosomes displayed long thin threads configuration, and thereafter homologous chromosomes partially paired at zygotene and fully synapsed at pachytene (Supplement Fig. S1A-C). At diplotene COs were formed between homologous chromosomes (Supplement Fig. S1D), and at diakinesis five pairs of bivalents occurred (Supplement Fig. S1E). At metaphase I, five pairs of bivalents were aligned at cell plate by bipolar pulling of spindles (Supplement Fig. S1F). Homologous chromosomes were then separated at anaphase I, and were temporally decondensed at interkinesis (Supplement Fig. S1G and H). In heat-stressed Col-0 plants, there was no obvious defect observed on leptotene and zygotene chromosomes (Supplement Fig. S1I and J); however, synapsis did not occur at pachytene and diplotene (Supplement Fig. S1K and L). Occurrence of ten univalents at diakinesis and metaphase I indicated that CO formation was abolished (Supplement Fig. S1M-Q). Fluorescence in situ hybridization (FISH) using a centromere-specific probe confirmed that the heat stress abolished bivalent formation (Supplement Fig. S2). In addition, we found that 83.33% diakinesis-staged pollen mother cells (PMCs) (*n* = 186) in heat-stressed plants displayed abnormal connections between univalents, which suggested that the high temperature induced interactions between non-homologous chromosomes (Supplement Fig. S1N-P, see red arrows). Heat-induced univalent formation leaded to unbalanced chromosome segregation at anaphase I and interkinesis (Supplement Fig. S1R-T), with resultant formation of aneuploid spores (Supplement Fig. S3A-D). Interestingly, in heat-stressed wild-type plants, we found that unicellular stage microspores occurred together with abnormal tetrad-staged meiocytes in the flower buds with the same size as control that only contained tetrads (Supplement Fig. S4). This implied that the high temperature may have influenced meiosis progression, either. Overall, these data demonstrated that high temperatures over fertility threshold completely suppress CO formation in Arabidopsis.

### Heat stress reduces SPO11-dependent DSB formation

Univalent formation under high temperatures phenocopied plants defective for DSB formation (Supplement Fig. S1M-Q) (De Muyt et al., 2007; Grelon et al., 2001; Huang et al., 2019; Stacey et al., 2006), we thus hypothesized that heat stress abolishes CO formation by interfering with DSB formation. We examined DSB number in heat-stressed wild-type and *spo11-1-1* plants by detecting γH2A.X, which marks phosphorylated H2A.X histone variants that specifically occur at DSB sites (Kurzbauer et al., 2012). Under control temperature, wild-type plants showed ~109 γH2A.X foci per meiocyte at zygotene stage, while the *spo11-1-1* mutant displayed a much lower count (~34) (Fig. 1A-C), indicating for defective DSB formation in the *spo11-1-1* plants. After 36-38°C treatment, we found that the number of γH2A.X foci in the wild-type plants was significantly lowered (~41) (Fig. 1A and D); however, γH2A.X foci in the *spo11-1-1* plants was not further reduced (~33) (Fig. 1A and E). We also checked the number of γH2A.X foci in wild-type plants stressed by 28°C and 32°C, the latter of which has been shown to induce univalent formation (De Storme and Geelen, 2020). The mild increase of temperature at 28°C did not influence the abundance of γH2A.X foci (~101), indicating for an unaffected DSB formation in line with the previous report (Supplement Fig. S5A-C) (Modliszewski et al., 2018). However, the 32°C-stressed plants showed significantly less γH2A.X dosage (~44) (Supplement Fig. S5A and D). To validate the measurement of γH2A.X foci, the secondary antibody was directly added to samples without immuno by the primary anti-γH2A.X antibody, and no γH2A.X foci was detectable (Supplement Fig. S6A and B). Taken together, these findings suggested that high temperature stress reduces SPO11-dependent DSB formation in Arabidopsis.

**Figure 1.**
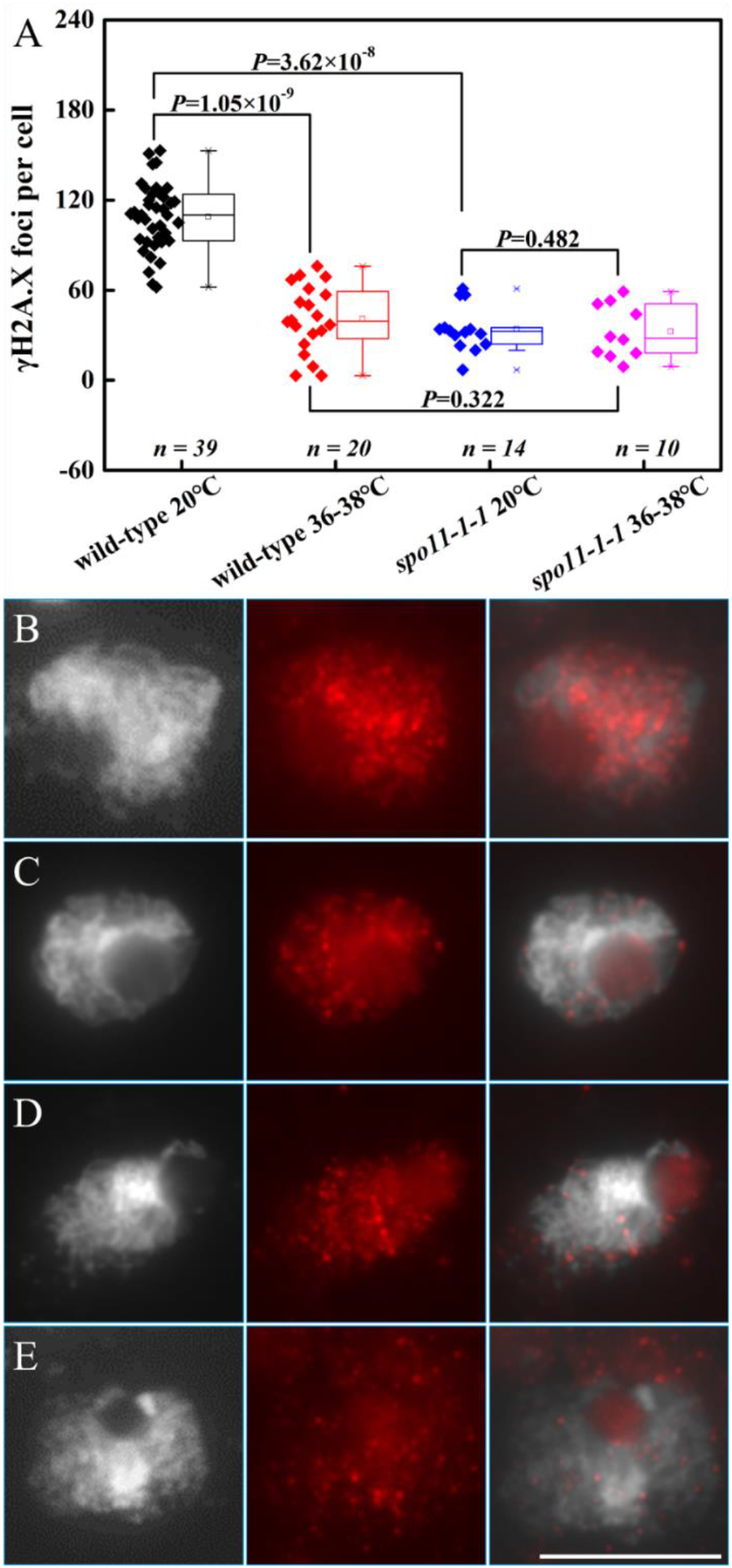
Immunolocalization of γH2A.X in heat-stressed wild-type Ws and *spo11-1-1* mutant plants. A, Graph showing the quantification of γH2A.X foci in wild-type and the *spo11-1-1* mutant under normal temperature and high temperature conditions. Two-side Mann-Whitney *U* test was performed. *n* indicates for the number of quantified cells. B and C, γH2A.X foci in wild-type (B) and *spo11-1-1* (C) plants under normal temperature conditions. D and E, γH2A.X foci in wild-type (D) and *spo11-1-1* (E) plants after heat stress. White: DAPI-stained chromosomes; Red: γH2A.X. Scale bar = 10 μm.

### Heat stress suppresses chromosome fragmentation in the *syn1* and *rad51* mutants

Loss of function of SYN1 or RAD51 causes chromosome fragmentation due to defective DSB repair in Arabidopsis (Bai et al., 1999; Lambing et al., 2020b; Su et al., 2017). The double *spo11 rad51* mutant rescues the chromosome fragmentation in *rad51* and phenocopies the singe *spo11* mutant (Hartung et al., 2007). To validate that heat stress reduces DSB formation, we analyzed meiosis I chromosome behaviors in the *syn1* and *rad51* mutants under heat stress. Under control temperature, numerous chromosome fragments were seen in diakinesis and metaphase I meiocytes in the *syn1* and *rad51* mutants (Fig. 2A and B, D and E). In anaphase I meiocytes of the mutants, considerable number of fragmented chromosomes were unbalancedly separated with some fragments remaining at the equator (Fig. 2C and F). After heat stress, we found that both the *syn1* and *rad51* mutants were able to produce ten univalents at diakinesis and metaphase I (Fig. 2G and I, M and O). At the same time, we observed that 89.16% (*n* = 83) and 87.88% (*n* = 33) diakinesis-staged meiocytes in the heat-stressed *syn1* and *rad51* plants, respectively, showed irregular interactions between univalents (Fig. 2H and N, see red arrows). Abnormal chromosome connections also occurred at metaphase I (Fig. 2J, P and Q). Heat-induced univalent formation leaded to unbalanced chromosome segregation at anaphase I in the mutants (Fig. 2K, L and R). Chromosome behaviors in the heat-stressed *syn1* and *rad51* plants mimicked the *spo11-1-1* mutant (Supplement Fig. S7), which indicated that heat stress suppresses chromosome fragmentation in *syn1* and *rad51*.

**Figure 2.**
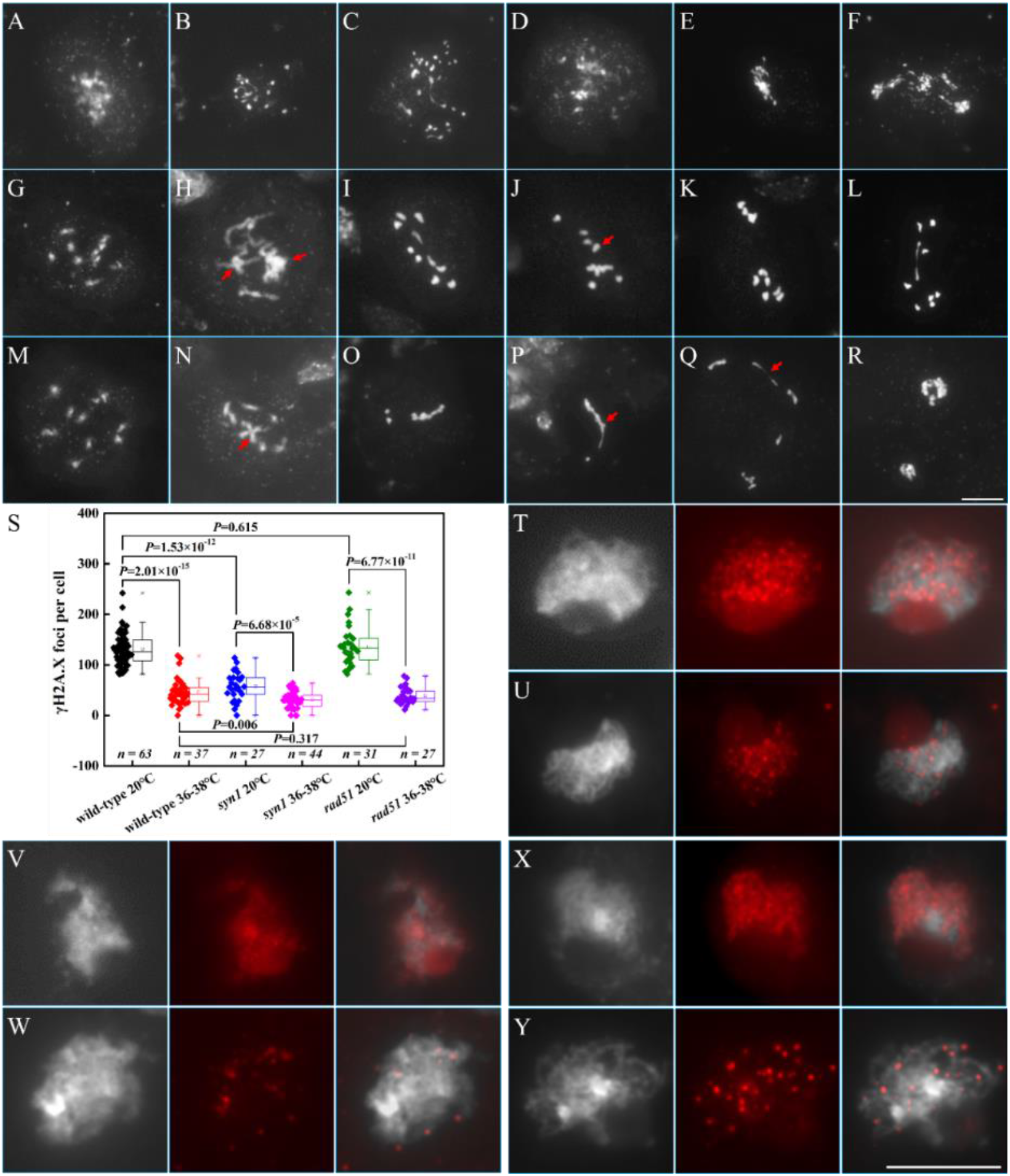
Heat stress partially suppresses chromosome fragmentation in the *syn1* and *rad51* mutants by lowering DSB formation. A-C, Diakinesis- (A), metaphase I- (B) and anaphase I-staged (C) PMCs in the *syn1* mutant under normal temperature conditions. D-F, Diakinesis- (D), metaphase I- (E) and anaphase I-staged (F) PMCs in the *rad51* mutant under normal temperature conditions. G-L, Diakinesis- (G and H), metaphase I- (I and J) and anaphase I-staged (K and L) PMCs in the *syn1* mutant after heat stress. M-R, Diakinesis- (M and N), metaphase I- (O-Q) and anaphase I-staged (R) PMCs in the *rad51* mutant after heat stress. Red arrows indicate irregular connections between univalent chromosomes. S, Graph showing the number of γH2A.X foci in heat-stressed wild-type, *syn1* and *rad51* plants. Two-side Mann-Whitney *U* test was performed. *n* indicates for the number of cells used for γH2A.X quantification. T and U, Immunolocalization of γH2A.X in wild-type under normal temperature conditions (T) and after heat stress (U). V and W, Immunolocalization of γH2A.X in the *syn1* mutant under normal temperature conditions (V) and after heat stress (W). X and Y, Immunolocalization of γH2A.X in the *rad51* mutant under normal temperature conditions (X) and after heat stress (Y). White: DAPI-stained chromosomes; Red: γH2A.X. Scale bars = 10 μm.

We next examined the γH2A.X number in the heat-stressed *syn1* and *rad51* mutants. Under control temperature, the *syn1* mutant showed significantly less γH2A.X number (~57) than the wild-type plants (~131) (Fig. 2S, T and V) as reported (Lambing et al., 2020b). After heat stress, the number of γH2A.X foci in the *syn1* mutant reduced further (~31), even less than the heat-stressed wild-type plants (~46) (Fig. 2S, U and W). This hinted that heat stress may reduce DSB formation independently of SYN1-mediated axis. On the other hand, the *rad51* mutant showed same levels of γH2A.X number as the wild-type plants under both the normal (~136) and high temperature conditions (~39) (Fig. 2S, X and Y). Taken together, these findings suggested that suppressed chromosome fragmentation in heat-stressed *syn1* and *rad51* plants was probably caused by lowered DSB formation.

### Heat stress reduces DMC1 foci on chromatin

DMC1 catalyzes MR-specific DSB repair, and in plants with impaired DSB formation, DMC1 is not detectable on chromatin (Da Ines et al., 2013; De Muyt et al., 2007; Kurzbauer et al., 2012; Sanchez-Moran et al., 2007). To further reveal that heat stress reduces DSB formation, we checked accumulation of DMC1 in heat-stressed wild-type plants. A novel anti-AtDMC1 antibody was generated, which showed specificity to AtDMC1 protein with no DMC1 foci detectable in the *atdmc1* mutant (Supplement Fig. S8). In wild-type plants incubated under control temperature, an average of 82 DMC1 foci per zygotene-staged meiocyte was recorded (Fig. 3A and B); while after heat stress, the number of DMC1 was significantly lowered (~18) (Fig. 3A and D), indicating that heat stress reduces the amount of DMC1 on chromatin. Since SYN1-mediated axis formation is required for normal DMC1 loading on chromatin (Lambing et al., 2020b), we then analyzed DMC1 dosage in *syn1* plants. Under control temperature, the *syn1* mutant showed significantly less DMC1 foci than wild-type plants (~56) (Fig. 3A and C); and the number decreased further (~13) (Fig. 3A and E) to a same level as in heat-stressed wild-type plants. These data suggested that heat stress reduces DMC1 foci on chromatin, and hinted that this impact may occur independently of the SYN1-mediated chromosome axis.

**Figure 3.**
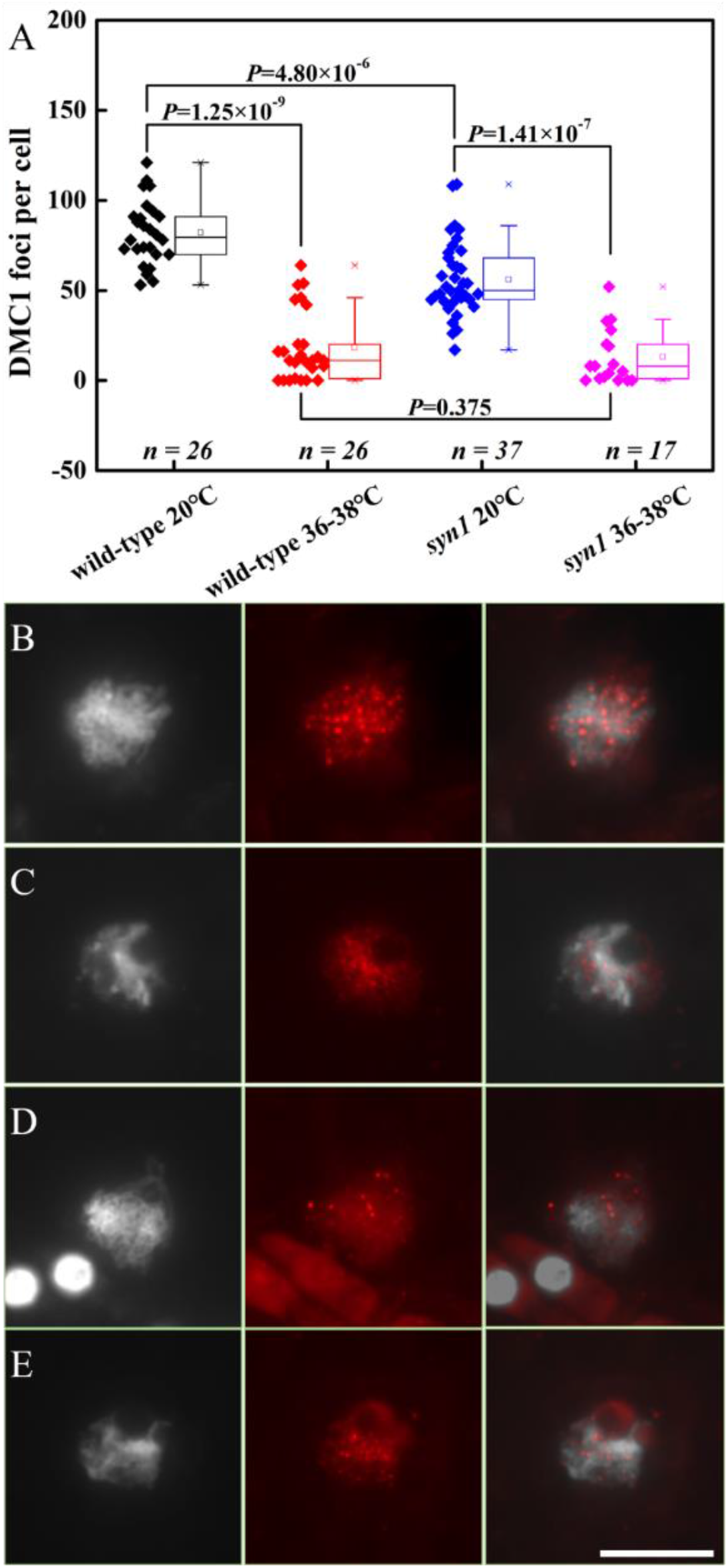
Immunolocalization of DMC1 in heat-stressed wild-type and *syn1* plants. A, Graph showing the number of DMC1 foci on zygotene chromosomes per PMC in wild-type and *syn1* plants under normal temperature and high temperature conditions. Two-side Mann-Whitney *U* test was performed. *n* indicates for the number of cells used for DMC1 foci quantification. B and C, DMC1 localization on zygotene chromosomes in wild-type (B) and *syn1* (C) plants under normal temperature conditions. D-E, DMC1 localization on zygotene chromosomes in wild-type (D) and *syn1* (E) plants after heat stress. White: DAPI-stained chromosomes; Red: DMC1. Scale bar = 10 μm.

### Heat stress does not affect SYN1-mediated chromosome axis

DSB formation relies on normal SYN1-mediated axis (Lambing et al., 2020b). To reveal whether heat-induced reduction of DSBs was owing to by impacted organization of SYN1-mediated axis, we performed immunostaining of SYN1 in both wild-type and the *spo11-1-1* plants. Under control temperature, wild-type plants showed linear SYN1 signals on zygotene and pachytene chromosomes, which was not detectable in the *syn1* mutant (Fig. 4A-C). Interestingly, we found that in heat-stressed wild-type plants, the loading of SYN1 on zygotene and pachytene chromosomes was not influenced (Fig. 4D and E), which indicated that the high temperature did not affect the formation of SYN1-mediated chromosome axis. Supportively, SYN1 accumulated normally on zygotene chromosomes in the *spo11-1-1* mutant under both control and high temperatures (Fig. 4F and G). Therefore, it is likely that heat stress does not affect SYN1, and reduces DSB formation independently of SYN1-mediated axis.

**Figure 4.**
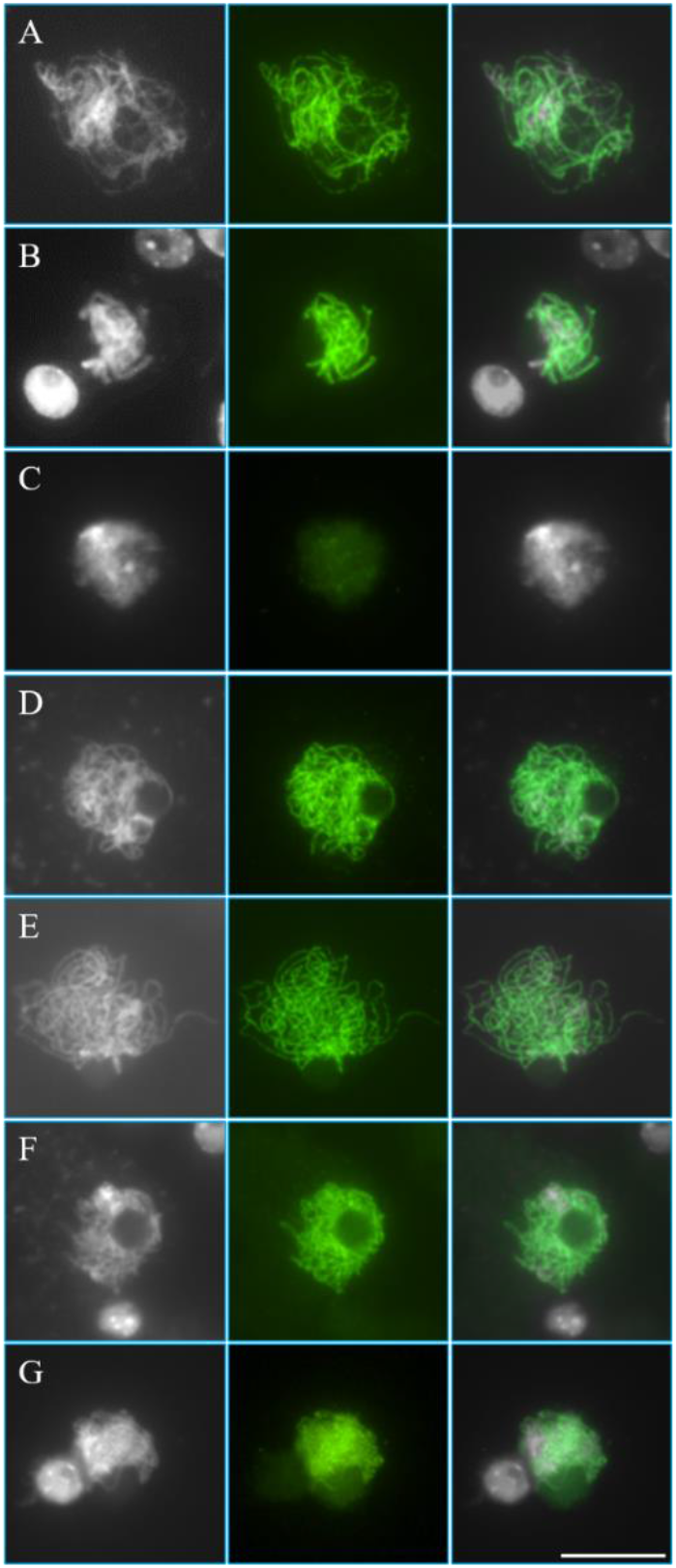
SYN1-mediated chromosome axis appears normal under heat stress. A and B, SYN1 immunolocalization on zygotene- (A) and pachytene-staged (B) chromosomes in wild-type plants under control temperature. C, SYN1 immunolocalization on zygotene-staged chromosome in the *syn1* mutant under control temperature. D and E, SYN1 immunolocalization on zygotene- (D) and pachytene-staged (E) chromosomes in wild-type plants after heat stress. F and G, SYN1 immunolocalization on zygotene-staged chromosomes in the *spo11-1-1* mutant under control temperature (F) and heat stress (G). White: DAPI-stained chromosomes; Green: SYN1. Scale bars = 10 μm.

### High temperatures affect ASY4-mediated chromosome axis

Axis formation requires functional and normal loading of ASY4 protein (Chambon et al., 2018). To determine whether heat stress influences chromosome axis, we generated an anti-AtASY4 antibody and examined ASY4 loading on prophase I chromosomes in wild-type and *syn1* plants. In wild-type plants grown under control temperature, numerous punctate ASY4 foci accumulated on leptotene chromosomes (Fig. 5A), which turned into linear configuration from early zygotene (Fig. 5B). In meiocytes from middle zygotene to late pachytene, ASY4 foci were fully associated with the entire chromosome regions (Fig. 5C-G). Sparse and punctate ASY4 foci occurred on diakinesis chromosomes indicating that ASY4 were unloaded after completion of MR (Fig. 5H). In the *syn1* mutant, which is defective for axis formation (Lambing et al., 2020b), ASY4 did not accumulate linearly on zygotene chromosomes, instead it showed incomplete and/or aggregated configurations (Supplement Fig. S9A and B). In wild-type plants stressed by 36-38°C, dotted ASY4 foci appeared on leptotene chromosomes and were sparser compared with control (Fig. 5I). At zygotene and/or pachytene, although there were some meiocytes showed normal ASY4 loading (Fig. 5K and M, zygotene; P, pachytene), about 56.92% (*n* = 65) meiocytes exhibited punctate ASY4 foci on chromatin (Fig. 5J, L, N, O, Q and R), indicating for an impacted ASY4 accumulation. At diakinesis, punctate ASY4 signals were associated with univalent chromosomes, whereas the number of dots seemed to be more than that of control (Fig. 5S and T). We observed similar ASY4 behaviors in wild-type plants stressed by 32°C (Fig. 5U-X). Taken together, these findings suggested that heat stress partially affects ASY4-mediated chromosome axis.

**Figure 5.**
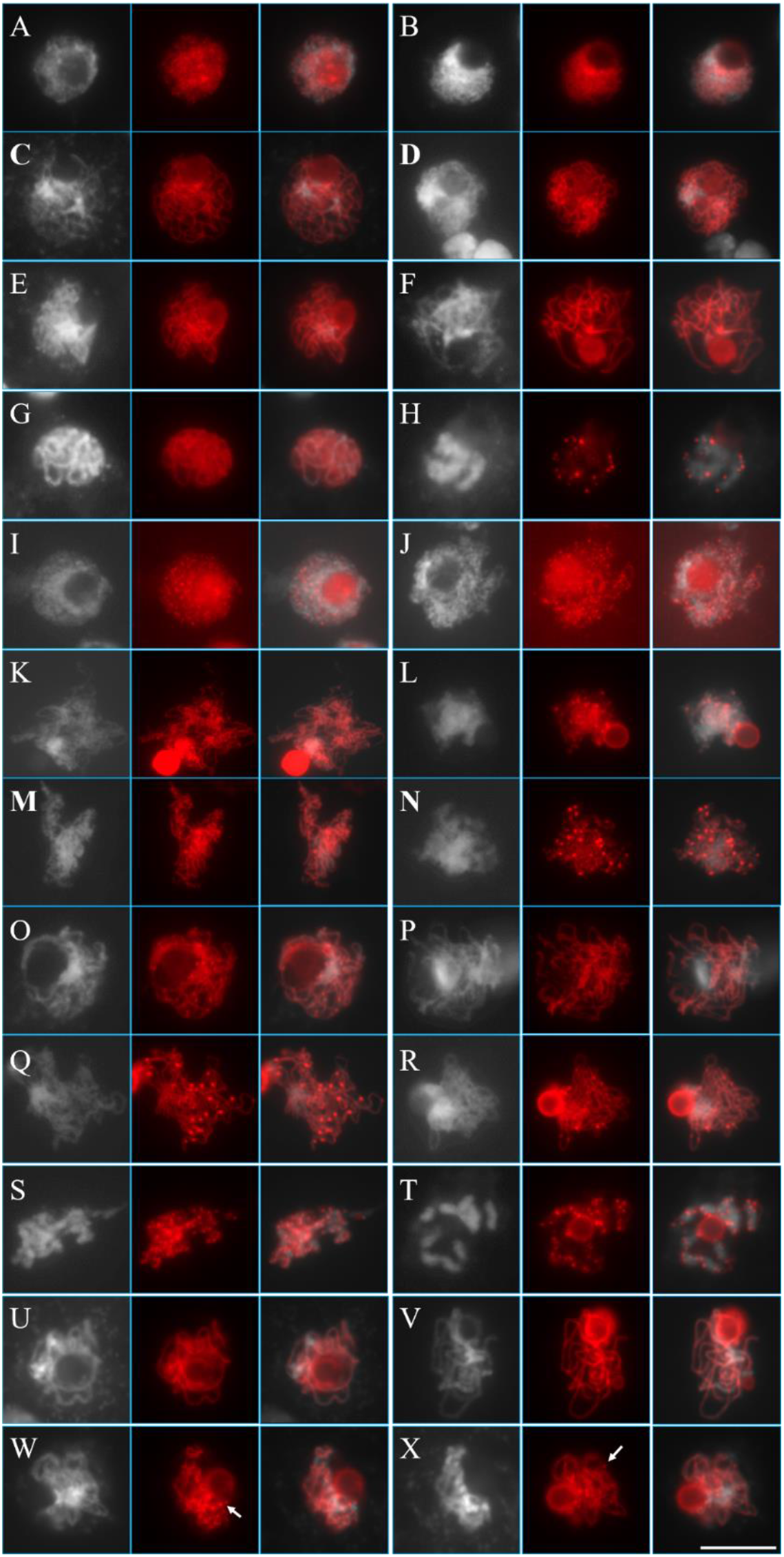
Heat stress induces destabilization of ASY4 on chromatin. A-H, Immunolocalization of ASY4 on leptotene (A), early zygotene (B), middle zygotene (C), late zygotene (D), early pachytene (E), middle pachytene (F), late pachytene (G) and diakinesis (H) chromosomes in wild-type plants under normal temperature conditions. I-T, Immunolocalization of ASY4 on leptotene (I), early zygotene (J), middle zygotene (K and L), late zygotene (M-O), middle pachytene (P and Q), late pachytene (R) and diakinesis (S and T) chromosomes in wild-type plants after 36-38°C heat stress. U-X, Immunolocalization of ASY4 on late zygotene (U and W) and middle pachytene (V and X) chromosomes in wild-type plants after 32°C heat stress. Write arrows indicate dotted ASY4 foci. White: DAPI-stained chromosomes; Red: ASY4. Scale bar = 10 μm.

### Heat stress affects ASY1-mediated lateral element of SC

Driven by the hypothesis that heat stress inhibits CO by affecting SC-dependent synapsis of homologous chromosomes, we examined lateral element of SC by immunostaining ASY1. In wild-type plants incubated under control temperature, punctate ASY1 foci occurred on leptotene chromosomes, which formed linear configuration from early zygotene (Fig. 6A and B). At middle zygotene, ASY1 foci were fully associated with the entire chromosomes at lateral axis regions (Fig. 6C). ASY1 foci were disassociated with some chromosome regions at late zygotene and early pachytene, when homologous chromosomes were almost fully paired (Fig. 6D and E). ASY1 disassociated further at middle pachytene when homologous chromosomes fully synapsed, and were completely unloaded at late pachytene (Fig. 6F and G). Remarkably, after heat stress, we observed that 49.66% (*n* = 147) meiocytes at leptotene, zygotene and/or pachytene showed sparse and punctate ASY1 foci (Fig. 6H and I, K-N), with other meiocytes showing normal ASY1 loading (Fig. 6J). Thus, these data suggested that heat stress induces instability or pre-mature destabilization at partial regions of ASY1-mediated lateral element of SC.

**Figure 6.**
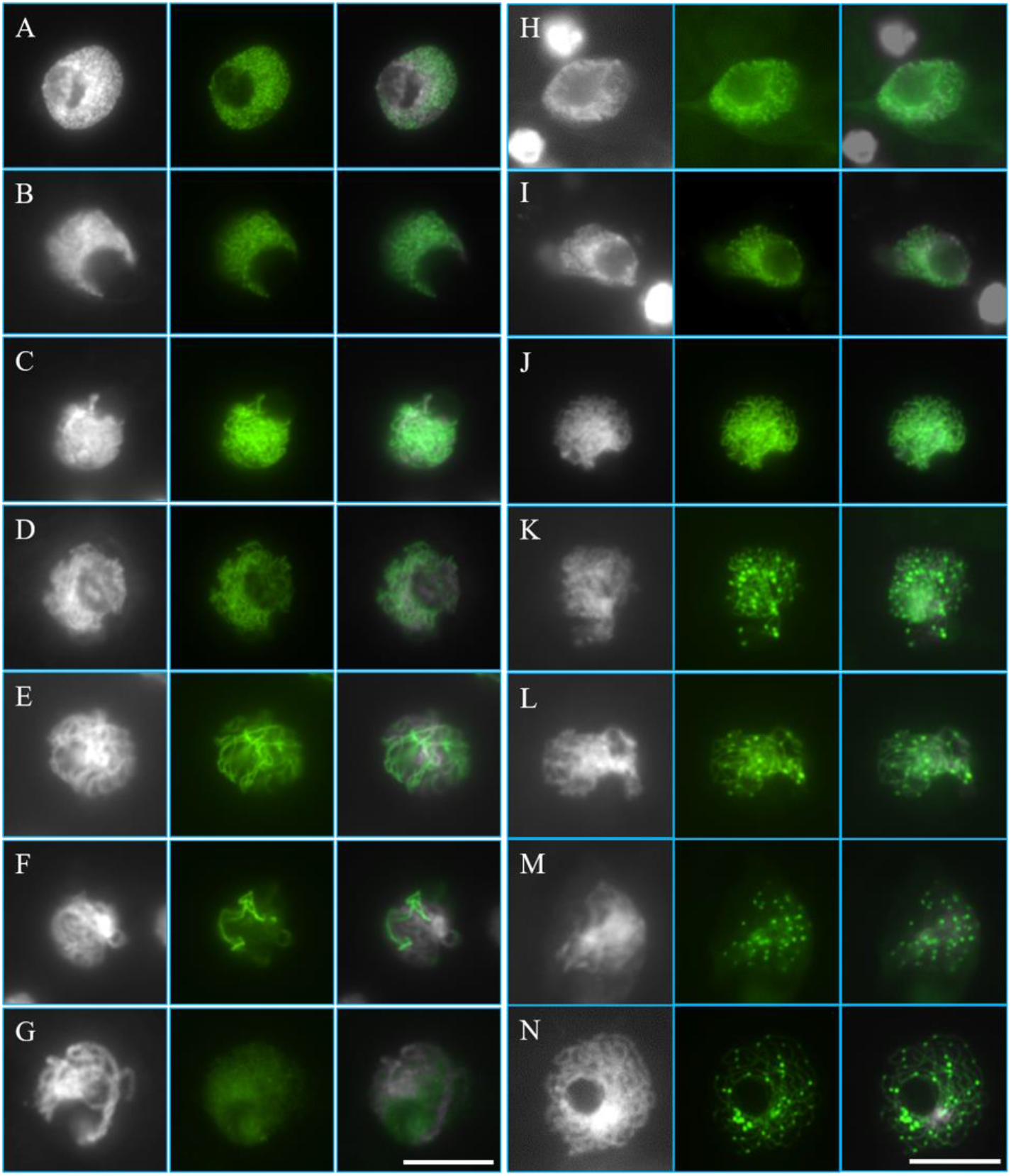
Immunolocalization of ASY1 in wild-type Col-0 plants under normal temperature and heat stress conditions. A-G, Chromosomes at leptotene (A), early zygotene (B), middle zygotene (C), late zygotene (D), early pachytene (E), middle pachytene (F) and late pachytene (G) under normal temperature conditions. H-N, Chromosomes at leptotene (H), early zygotene (I), middle zygotene (J-L), late zygotene (M) and middle pachytene (N) after heat stress. White: DAPI-stained chromosomes; Green: ASY1. Scale bars = 10 μm.

### Heat-induced ASY1 abnormity relies on the presence of SYN1 but not DSB formation

To understand whether heat-induced alterations in ASY1 loading was associated with the impact on DSB formation, we analyzed ASY1 in heat-stressed *spo11-1-1* plants. Under control temperature, ASY1 accumulated normally on zygotene chromosomes in *spo11-1-1* as in wild-type plants (Fig. 7A and B). After heat stress, we observed punctate ASY1 foci in both the wild-type and *spo11-1-1* plants (Fig. 7D and E). These suggested that the organization of ASY1-mediated lateral element of SC and its response to heat stress does not rely on DSB formation. Since building of ASY1-mediated lateral element of SC relies on functional SYN1 (Lambing et al., 2020b), we then checked whether heat-interfered loading of ASY1 was SYN1-dependent. Interestingly, we found that the *syn1* mutant incubated under both 20°C and 36-38°C showed same ASY1 configuration; i.e. incomplete or fragmented ASY1 loading on zygotene chromosomes (Fig. 7C and F), which thus hinted that heat stress influences ASY1-mediated lateral element of SC relying on the presence of SYN1.

**Figure 7.**
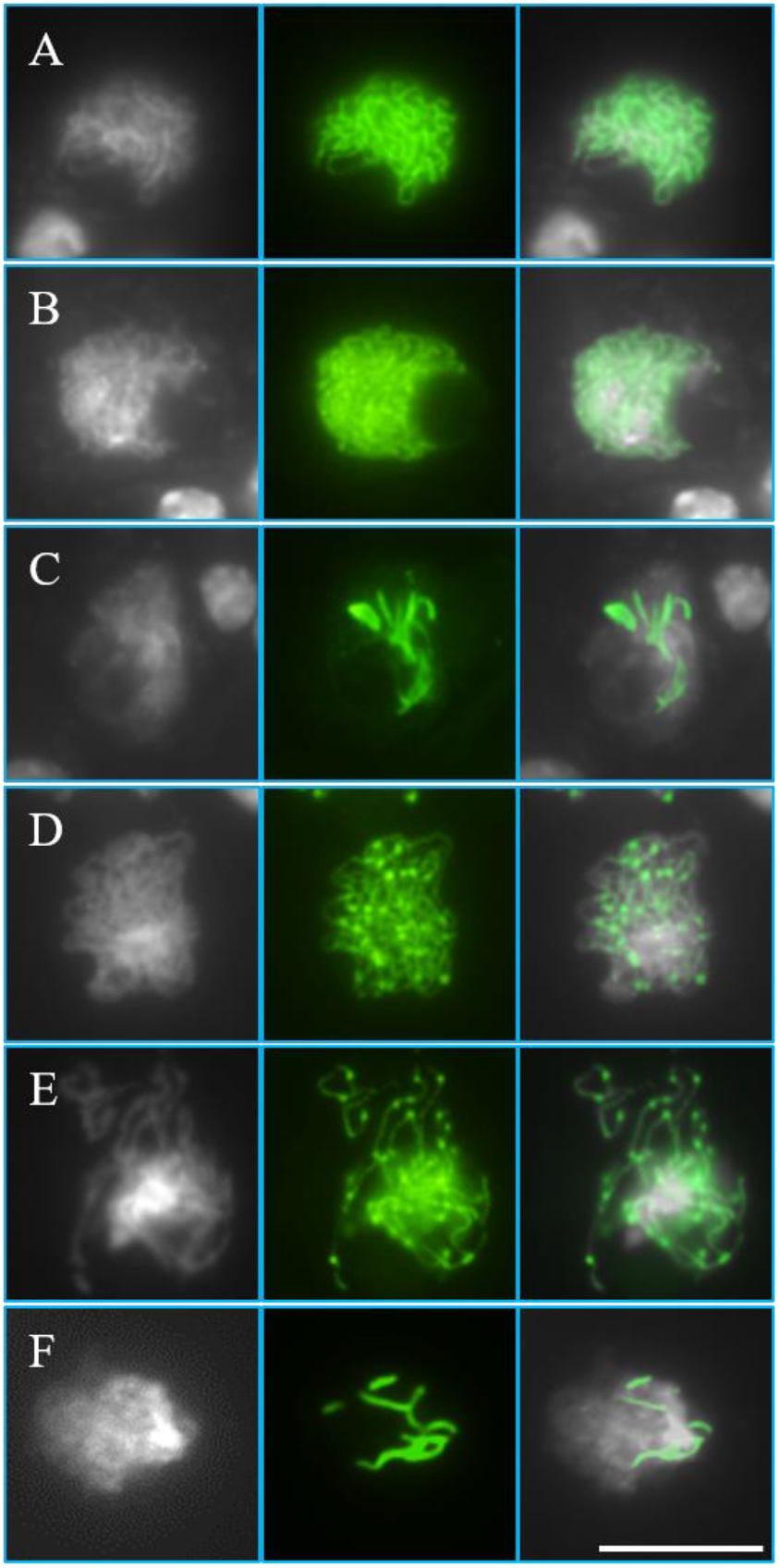
Immunolocalization of ASY1 in heat-stressed wild-type, *spo11-1-1* and *syn1* plants. A-C, ASY1 localization on zygotene chromosomes of wild-type (A), *spo11-1-1* (B) and *syn1* mutants (C) under normal temperature conditions. D-F, ASY1 localization on zygotene chromosomes of heat-stressed wild-type plants (D), and the *spo11-1-1* (E) and *syn1* mutants (F). White: DAPI-stained chromosomes; Green: ASY1. Scale bar = 10 μm.

### Heat stress disrupts central element of SC

Central element of SC was examined by performing immunolocalization of ZYP1. In control plants, ZYP1 linearly accumulated on zygotene chromosomes, and were fully associated with the central regions of synapsed homologous chromosomes at pachytene (Fig. 8A and D). In heat-stressed plants, however, ZYP1 signals displayed dotted, aggregated and/or fragmented configurations on zygotene and/or pachytene chromosomes (Fig. 8B and C, E and F). This finding indicated that heat stress disrupts the formation of ZYP1-based central element of SC. ZYP1-dependent homology synapsis requires normal DSB formation (De Muyt et al., 2007; Grelon et al., 2001; Stacey et al., 2006), which drove us to check whether heat-induced defective ZYP1 was caused by the suppressed DSB formation. Under control temperature, 94.74% zygotene and/or pachytene meiocytes in the *spo11-1-1* mutant displayed small and dotted, or aggregated ZYP1 foci (Fig. 8G and H), while the other 5.26% meiocytes exhibited empty ZYP1 loading (Fig. 8I, *n* = 19). In contrast, 86.67% zygotene and/or pachytene meiocytes in heat-stressed *spo11-1-1* plants had no ZYP1 foci (Fig. 8J, *n* = 15). The largely increased ratio of meiocytes with completely failed ZYP1 loading in *spo11-1-1* suggested that heat stress may directly influence ZYP1-dependent SC assembly.

**Figure 8.**
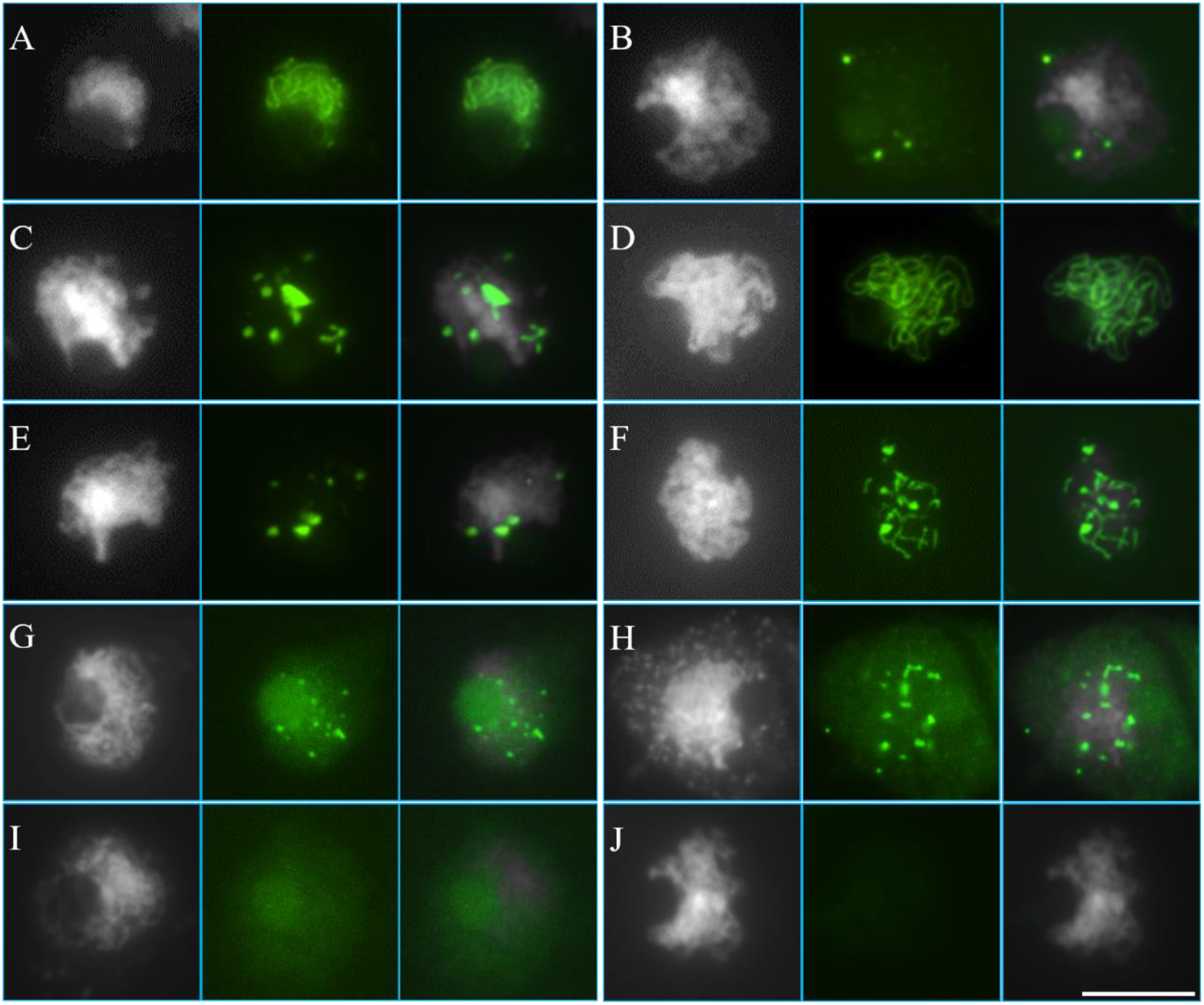
Immunolocalization of ZYP1 in heat-stressed wild-type and *spo11-1-1* mutant plants. A-C, ZYP1 localization on zygotene chromosomes of wild-type under normal temperature (A) and heat stress (B and C) conditions. D-F, ZYP1 localization on pachytene chromosomes of wild-type under normal temperature (D) and heat stress (E and F) conditions. G-J, ZYP1 localization on *spo11-1-1* mutant chromosomes under normal (G-I) and high temperature (J) conditions. White: DAPI-stained chromosomes; Green: ZYP1. Scale bar = 10 μm.

### Expression pattern of MR-related genes under heat stress

We performed quantitative RT-PCR (qRT-PCR) to address whether heat stress influences MR by impacting the expression of MR-related genes. Arabidopsis heat shock sensor *HSFA1a* was used as positive control (Yoshida et al., 2011), which showed 1.71-fold increased expression level under heat stress (Fig. 9A, *P* < 0.05). Interestingly, the expression of *SPO11-1*, *PRD1*, *2* and *3*, which are required for DSB formation, were not significantly altered by the heat stress (Fig. 9B-E, *P* > 0.05). This implied that high temperature may suppress DSB formation not by lowering the transcription of DSB catalyzers. Two phosphatidylinositol 3 kinase-like (PI3K) protein kinases ATR and RAD3-Related (ATR) and Ataxia-Telangiectasia Mutated (ATM), which phosphorylate histone variant H2AX at DSB sites and evoke DSB repair (Amiard et al., 2011), were not influenced and upregulated, respectively (Fig. 9F, *P* > 0.05; G, *P* < 0.01). In addition, we found that *RAD51* was significantly downregulated by the high temperature, while *DMC1* was not changed (Fig. 9H, 0.56-fold, *P* < 0.001; I, *P* > 0.05). Remarkably, chromosome axis factor *SYN1* was upregulated; on the contrary, *ASY3* and *ASY4* were largely downregulated by the heat stress (Fig. 9J, 2.44-fold, *P* < 0.05; K, 0.53-fold, *P* < 0.001; L, 0.44-fold, *P* < 0.001). This suggested that heat stress may affect ASY4-mediated chromosome axis via an impacted transcription of axis factors. Moreover, SC components *ASY1* and *ZYP1a* displayed the same expression level as control (Fig. 9M and N, *P* > 0.05). The expression of *OSD1* and *TAM*, which control meiotic cell cycle transition in Arabidopsis (d’Erfurth et al., 2010; d’Erfurth et al., 2009), were not changed and increased under heat stress, respectively (Fig. 9O, *P* > 0.05; P, 1.47-fold, *P* < 0.05). The increased expression of *TAM* hinted that the high temperature may promote transition of meiotic cell cycles and accelerate meiosis progression (Supplement Fig. S4).

**Figure 9.**
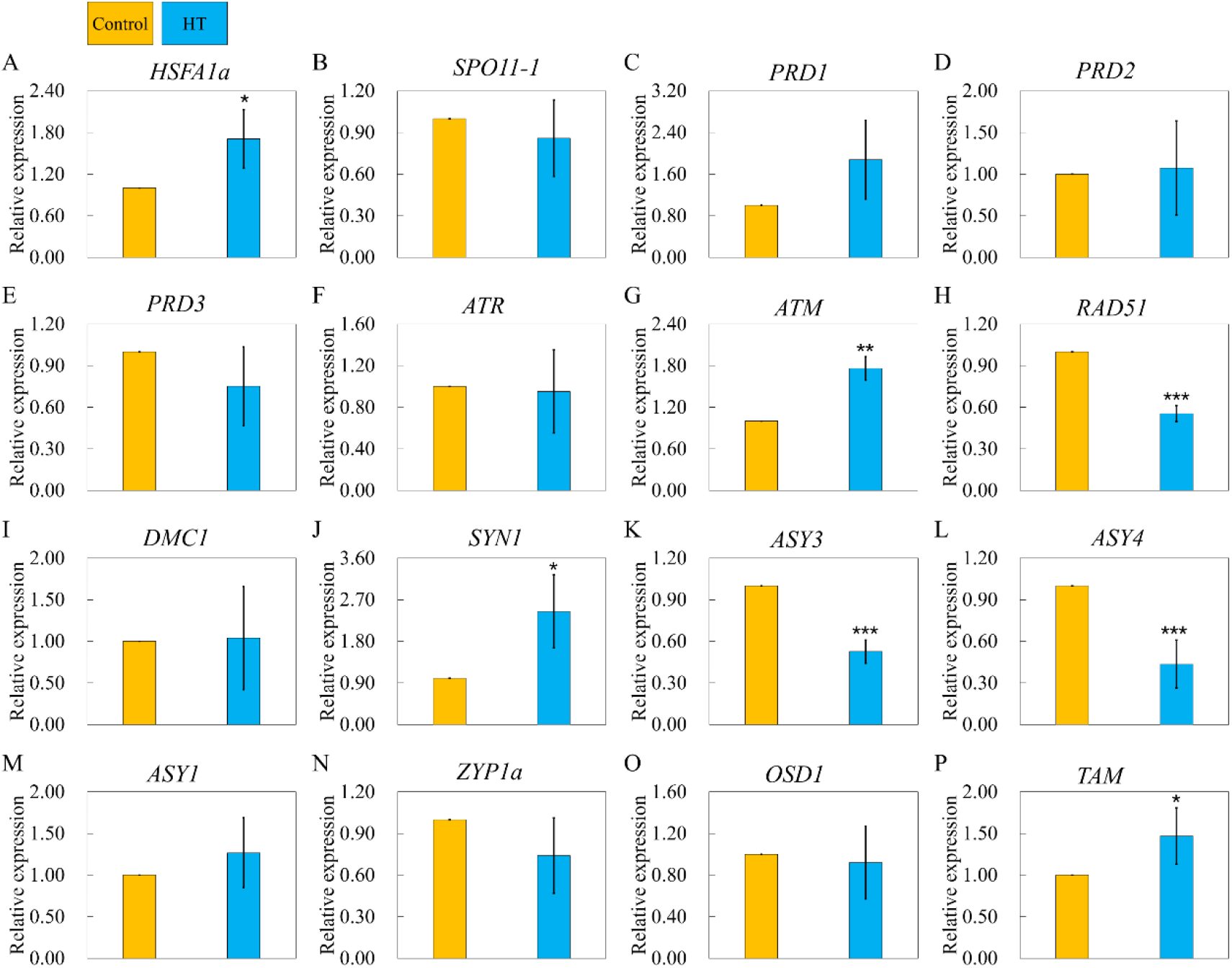
Expression pattern of meiosis-related genes under heat stresses. A-P, Relative expression fold changes of *HSFA1a* (A), *SPO11-1* (B), *PRD1* (C), *PRD2* (D), *PRD3* (E), *ATR* (F), *ATM* (G), *RAD51* (H), *DMC1* (I), *SYN1* (J), *ASY3* (K), *ASY4* (L), *ASY1* (M), *ZYP1a* (N), *OSD1* (O) and *TAM* (P) after heat stress. Gold columns indicate control temperature and blue columns indicate high temperature conditions, respectively. Significance level was set as *P* < 0.05. * indicates for *P* < 0.05, ** indicates for *P* < 0.01, *** indicates for *P* < 0.001.

## Discussion

### Heat stress suppresses MR via compromised DSB formation

MR in plants is sensitive to variations of environmental temperature (Bomblies et al., 2015; Liu et al., 2019; Modliszewski and Copenhaver, 2017). Lloyd et al. proposed that MR rate of Arabidopsis under different temperature conditions within fertility-tolerable range displays a ‘U-shape’ pattern, in which both decreased (8°C) and increased (28°C) temperatures promote MR rate (Lloyd et al., 2018). A similar influential pattern was found in wheat incubated between 10~26°C (Coulton et al., 2020), which suggests that the positive impact of higher temperatures within fertility threshold on MR frequency may be conserved between dicots and monocots. Under a higher temperature at 32°C, Arabidopsis plants produce univalents indicating for a lowered MR frequency (De Storme and Geelen, 2020). Here, we showed that extreme high temperatures over fertility-tolerable threshold (36-38°C) fully inhibits CO formation in Arabidopsis. Therefore, it is likely that in Arabidopsis, MR rate reaches peak between temperature interval from 28 to 32°C.

Moderate elevation of temperature promotes MR in Arabidopsis via enhanced activation of type-I CO formation pathway but not by influencing DSB formation (Lloyd et al., 2018; Modliszewski et al., 2018). Similarly, in *Schizosaccharomyces pombe*, MR rate is moderately changed without alteration in DSB formation under temperatures within fertile range from 16 to 33°C (Brown et al., 2019). It thus is possible that DSB catalyzation system has a threshold to increase of environmental temperature, below which the system functions normally and generates DSBs. Here, we showed that in Arabidopsis, DSB formation is significantly reduced at and above 32°C (Fig. 1A and D; Supplement Fig. S5D). In support, the *syn1* and *rad51* mutants stressed by 36-38°C have lowered DSB dosage, and are able to produce ten univalents, which phenocopies *spo11-1-1* plants (Fig. 2; Supplement Fig. S7). In *S. cerevisiae*, it has been reported that DSB repair defect (manifested by calculated gene conversion rate) in the *rad55* and/or *rad57* mutants is more severe at lower temperatures (Pohl and Nickoloff, 2008). These facts imply that suppression of DSB formation under high temperatures may be a conserved phenomenon among eukaryotes. We found that the expression of *SPO11-1*, *PRD1*, *PRD2* and *PRD3*, which are required for DSB formation (Carballo et al., 2008; De Muyt et al., 2009; De Muyt et al., 2007; Grelon et al., 2001; Stacey et al., 2006), are not changed under heat stress (Fig. 9B-E). Therefore, it is likely that heat stress reduces DSB formation not by lowering the transcription of the corresponding DSB catalyzers.

Since CO formation and MR are suppressed in Arabidopsis plants defective for DSB generation (De Muyt et al., 2007; Grelon et al., 2001; Hartung et al., 2007; Stacey et al., 2006), our study reveals that beyond a tolerable temperature range, high temperatures interfere with MR by reducing SPO11-dependent DSB formation in Arabidopsis. There hence may exist a homeostasis between a positive impact of high temperature on type-I CO formation machineries and a negative influence on DSB formation in Arabidopsis: a limited temperature elevation within the tolerable range of DSB formation (e.g. 28°C) promotes the activation of ZMM proteins and increases type-I CO frequency. However, when the temperature reaches a point where SPO11-dependent DSB formation is interfered (e.g. from 32°C), CO rate decreases and is fully inhibited at 36-38°C.

### Chromosome axis and SC assembly are prominent targets of high temperatures

A recent study in Arabidopsis have revealed that at 32°C, ASY1 foci appear normal but linear configuration of ZYP1 on pachytene chromosomes is interrupted (De Storme and Geelen, 2020). In barley, a moderate high temperature causes a slight but significant decrease of CO formation by affecting ZYP1-dependent SC assembly (Higgins et al., 2012). Here, we showed that accumulation of ASY4 on chromatin is influenced under high temperatures at 32 and 36-38°C (Fig. 5). At the same time, we showed that ASY1-associated lateral element of SC is partially influenced (Fig. 6), and the ZYP1-dependent central element of SC is severely disrupted under 36-38°C (Fig. 8). These facts indicate that high temperatures have a predominant impact on the assembly and/or stability of chromosome axis and SC (Morgan et al., 2017). Considering the essential requirement of axis and SC to successful CO formation (Ferdous et al., 2012; Higgins et al., 2005; Sanchez-Moran et al., 2007), heat-induced suppression of MR could at least partially result from an interfered axis and SC formation.

We did not find any meiocyte with defective SYN1 loading under high temperature, suggesting that SYN1-mediated axis formation is not affected by heat stress (Fig. 4D and E). Thus, the observed defects in ASY4, ASY1 and ZYP1 configuration are not likely caused by alteration in SYN1. Nevertheless, we found that heat stress does not induce dotted ASY1 foci in the *syn1* mutant (Fig. 7C and F), suggesting that heat stress disturbs the assembly of ASY1 relying on the presence of SYN1. Molecular insights are needed to understand the role of SYN1 in mediating the response of ASY1 to high temperature. The expression of *ASY1* and *ZYP1* is not changed under high temperature, but *ASY3* and *ASY4* do (Fig. K-M). This suggests that heat stress may affect axis formation via modulated transcription of axis components, but influence SC by other mechanisms; e.g. ASY1 affinity to chromatin and pairing of homologous chromosomes. Indeed, heat stress can directly interfere with MR by influencing the activity of proteins involved in DSB processing and/or CO formation (Börner et al., 2004; Hotta et al., 1988; Smith et al., 2003). On the other hand, we observed similar alterations in loading of ASY1 and ASY4 on chromosomes under heat stress (Fig. 5 and Fig. 6), which suggested that heat induces instability of ASY1-mediated lateral element of SC and ASY4-mediated axis formation through a same signaling pathway. ASY1 loading on chromatin requires the presence and function of ASY3, which in turn relies on ASY4 (Chambon et al., 2018; Ferdous et al., 2012; Lambing et al., 2020a; Yang et al., 2019). It thus is possible that accumulation of ASY3 on chromatin is also interfered, and heat-induced destabilization of ASY1 is a downstream event of the affected ASY4.

The *asy1* mutant has normal DSB formation and axis structuring, but has no homolog synapsis (Ferdous et al., 2012; Sanchez-Moran et al., 2007). So, heat-induced DSB reduction is not likely to be caused by the disorganized ASY1-associated axis. In addition, since we can find 50% meiocytes with normal ASY1 loading, but did not see any meiocyte with normal ZYP1 accumulation under heat stress, the impaired ZYP1-dependent homolog synapsis could not be mainly owing to the interfered ASY1 function. However, DSB formation and ZYP1-dependent homolog synapsis relies on ASY3-medaited axis formation (Chambon et al., 2018; Ferdous et al., 2012; Sanchez-Moran et al., 2007), therefore, the compromised DSB formation and synapsis of homologous chromosomes could result from the impacted chromosome axis formation.

### DMC1-dependent MR is a potential target of heat stress

In wheat, high temperature lowers CO formation, and this effect is more pronounced in the *tadmc1* mutant (Draeger et al., 2019). In our study, we showed that the number of DMC1 foci decreases under heat stress (Fig. 3), which supports the notion that DMC1 is a potential target of high temperature (Brown et al., 2019). The expression of *DMC1* under heat stress is not significantly changed, but *RAD51* is lowered (Fig. 9I). It is possible that heat reduces DMC1 accumulation by influencing its affinity to chromatin via impacted RAD51 (Cloud et al., 2012; Kurzbauer et al., 2012). It is also possible that the reduced DMC1 foci is a downstream effect of compromised DSB formation, and/or the destabilized chromosome axis (De Muyt et al., 2007; Ferdous et al., 2012; Sanchez-Moran et al., 2007).

## Supplemental Files

The following files are available in the online version of this article.

Supplement Figure S1. DAPI-staining of meiotic chromosomes in heat-stressed wild-type Col-0 plants.

Supplement Figure S2. FISH analysis of chromosome dynamics in heat-stressed wild-type Col-0 plants.

Supplement Figure S3. Heat stress induces abnormal tetrad-staged PMCs.

Supplement Figure S4. Meiotic products of heat-stressed wild-type Col-0 plants stained by orcein.

Supplement Figure S5. Quantification of γH2A.X in wild-type Col-0 plants incubated at 20°C, 28°C and 32°C conditions.

Supplement Figure S6. Specificity test of anti-γH2A.X antibody.

Supplement Figure S7. Meiotic spread in the *spo11-1-1* mutant under control temperature. Supplement Figure S8. Immunolocalization of DMC1 in the *dmc1* mutant.

Supplement Figure S9. Immunolocalization of ASY4 in the *syn1* mutant under normal temperature conditions.

Supplement Table S1. Primers used in this study.

## Material and Methods

### Plant materials and growth conditions

*Arabidopsis thaliana* ecotypes Columbia-0 (Col-0) and Wassilewskija (Ws) plants were used as wild-type plants. The *syn1-1* (SALK_137095) (Ws background), *rad51* (SAIL_873_C08) (Col-0 background), *spo11-1-1* (Ws background) (Grelon et al., 2001) and *dmc1* (SALK_056177) (Col-0 background) (Sanchez-Moran et al., 2007) mutants were used in the study. Seeds were germinated in soil for 6-8 days and seedlings were transferred to soil and cultivated in growth chambers with a 16 h day/8 h night, 20°C, and 50% humidity condition. For temperature stress treatment, young flowering plants were transferred to a humid chamber with a 16 h day/8 h night and at 28, 32 and/or 36-38°C conditions for 24 h, respectively. All the treatment assays started from 8:00-10:00 AM. Meiosis-staged flower buds were fixed right away upon the finish of treatment.

### Cytology and quantification of γH2A.X and DMC1 foci

Orcein staining of meiotic products was performed by referring to (Lei et al., 2020). For the quantification of fluorescent foci of γH2A.X and DMC1, we referred to (Xue et al., 2018; Yao et al., 2020). Images of DAPI-stained chromosome signals and RFP-channeled protein foci signals were merged, and only the foci merged onto chromosomes were counted. Foci were identified manually and counted automatically by the Image J count tool.

### Generation of antibodies

Anti-AtZYP1 (rabbit) and AtDMC1 (rabbit) antibodies were generated by referring to (Yu et al., 2012); anti-AtASY4 peptide antisera was raised in rabbits against the amino acid sequence DSVNKSARGKMLQLKM of AtASY4 conjugated to KLH, by GL Biochem, Shanghai, Ltd: www.glschina.com. Anti-AtASY1 (rabbit) peptide antisera was raised in rabbits against the amino acid sequence NCSQASQDRRGRKTS of AtASY1 conjugated to KLH. Purification of antibodies was performed using polypeptide affinity followed by Elisa test.

### Immunolocalization and fluorescence in situ hybridization (FISH)

Immunolocalization assays were performed by referring to (Chelysheva et al., 2010; Wang et al., 2014). Antibodies against ASY1 (rabbit), ZYP1 (rabbit), DMC1 (rabbit), and γH2A.X (rabbit) were diluted by 1:100; antibody against AtASY4 (rabbit) was diluted by 1:200; antibody against SYN1 (rabbit) (Wang et al., 2020) was diluted by 1:500. The secondary antibodies, i.e. Goat anti-Rabbit IgG (H+L) Cross-Adsorbed Secondary Antibody Alexa Fluor 555 (Invitrogen, A32732) and Goat anti-Rabbit IgG (H+L) Highly Cross-Adsorbed Secondary Antibody Alexa Fluor Plus 488 (Invitrogen, A32731) were diluted by 1:400. FISH and used centromere-specific probes were referred to (Lei et al., 2020).

### Quantitative RT-PCR (qRT-PCR)

Total RNA was isolated from meiosis-staged flower buds of Arabidopsis Col-0 plants grown under normal temperature conditions (20°C) and after 36-38°C treatment for 24 h, respectively. First-strand cDNA was synthesized using the PrimeScriptTM II 1st Strand cDNA Synthesis Kit (TAKARA, Kusatsu, Japan) according to the manufacturer’s protocol. qRT-PCR was performed on a QuantStudio 1 Real-Time PCR System (Applied BiosystemsTM, Thermo Fisher Scientific) using the Applied BiosystemsTM PowerUpTM SYBRTM Green Master Mix (Thermo Fisher Scientific, Vilnius, Lithuania). The reaction mix was prepared with 10.0 μL PowerUpTM SYBRTM Green Master Mix, 0.4 μL 10 μM forward primer, 0.4 μL 10 μM reverse primer, 75 ng cDNA, and nuclease-free water up to a total volume by 20.0 μL. For both the control and heat stress conditions, three biological replicates were analyzed, and for each sample, three technical replicates for each targeted gene were performed. *EF1αA4* (At5g60390) was used as the reference gene (Parra-Rojas et al., 2019), and the relative expression fold-change was calculated using the Comparative CT method. Significance analysis was performed using the one-way ANOVA test, and significance level was set as 0.05. The used primers are listed in Supplement Table S1.

### Microscopy

Fluorescence images were recorded using an Olympus IX83 inverted fluorescence microscope with a X-Cite lamp and a Prime BSI camera. Bifluorescent images and Z-stacks were processed using Image J. Brightness and contrast setting of pictures were adjusted using PowerPoint 2016.

## Accession Numbers

Accession numbers of all genes studied in this work are: *SPO11-1* (AT3G13170), *ASY1* (AT1G67370), *ASY3* (AT2G46980), *ASY4* (AT2G33793), *ZYP1a* (AT1G22260), *SYN1* (AT5G05490), *DMC1* (AT3G22880), *RAD51* (AT5G20850), *ATM* (AT3G48190), *ATR* (AT5G40820), *HSFA1a* (AT4G17750) and *EF1αA4* (At5g60390).

## Acknowledgement

We thank very much for the suggestions and comments from Dr. Wojtek Pawlowski (Cornell University) for this work. We appreciate Dr. Yingxiang Wang (Fudan University) for kindly sharing the *syn1*, *rad51*, *spo11-1-1* and *dmc1* mutants, and the anti-SYN1 antibody. We thank Dr. Yan He (China Agricultural University) very much for kindly sharing the anti-γH2A.X antibody. We thank Huiqi Fu and Xiaohong Zhang (South-Central University for Nationalities) for their help on tetrad analysis, and we thank Dr. Cong Wang (Fudan University) and Dr. Wenqing Shi (Institute of Genetics and Developmental Biology, Chinese Academy of Sciences) for the technical help on immunolocalization assays. Especially, we thank Mrs. Meirong Huang, Mrs. Manxiang Zhu, Mr. Donglin Liu, Mrs. Jiajia Zhao and Ms. Zipei Liu for the support during the occurrence of coronavirus COVID-19 global cases.

