## Supplementary material for "Heat Stress Interferes with Formation of Double-Strand Breaks and Homolog Synapsis in *Arabidopsis thaliana*": Supplenent File

[1]College of Life Sciences, South-Central University for Nationalities, Wuhan 430074, China;
[2]The Modernization Engineering Technology Research Center of Ethnic Minority Medicine of Hubei Province, School of Pharmaceutical Sciences, South-Central University for Nationalities, Wuhan 430074, China;
[3]College of Life Sciences, Shanghai Normal University, Shanghai 200234, China;
[4]School of Integrative Plant Science, Cornell University, Ithaca, NY 14853, United States of America;
[5]College of Life Science and Technology, Huazhong Agricultural University, Wuhan, 430070;
[6]Key Laboratory of Horticultural Plant Biology (Ministry of Education), Huazhong Agricultural University, Wuhan, 430070, China;
[7]Agricultural Biochemistry Department, Faculty of Agriculture, Zagazig University, 44511 Zagazig, Egypt;
[#]Authors with equal contribution;

**Supplemental Figures**

*Supplemental figure S1.*

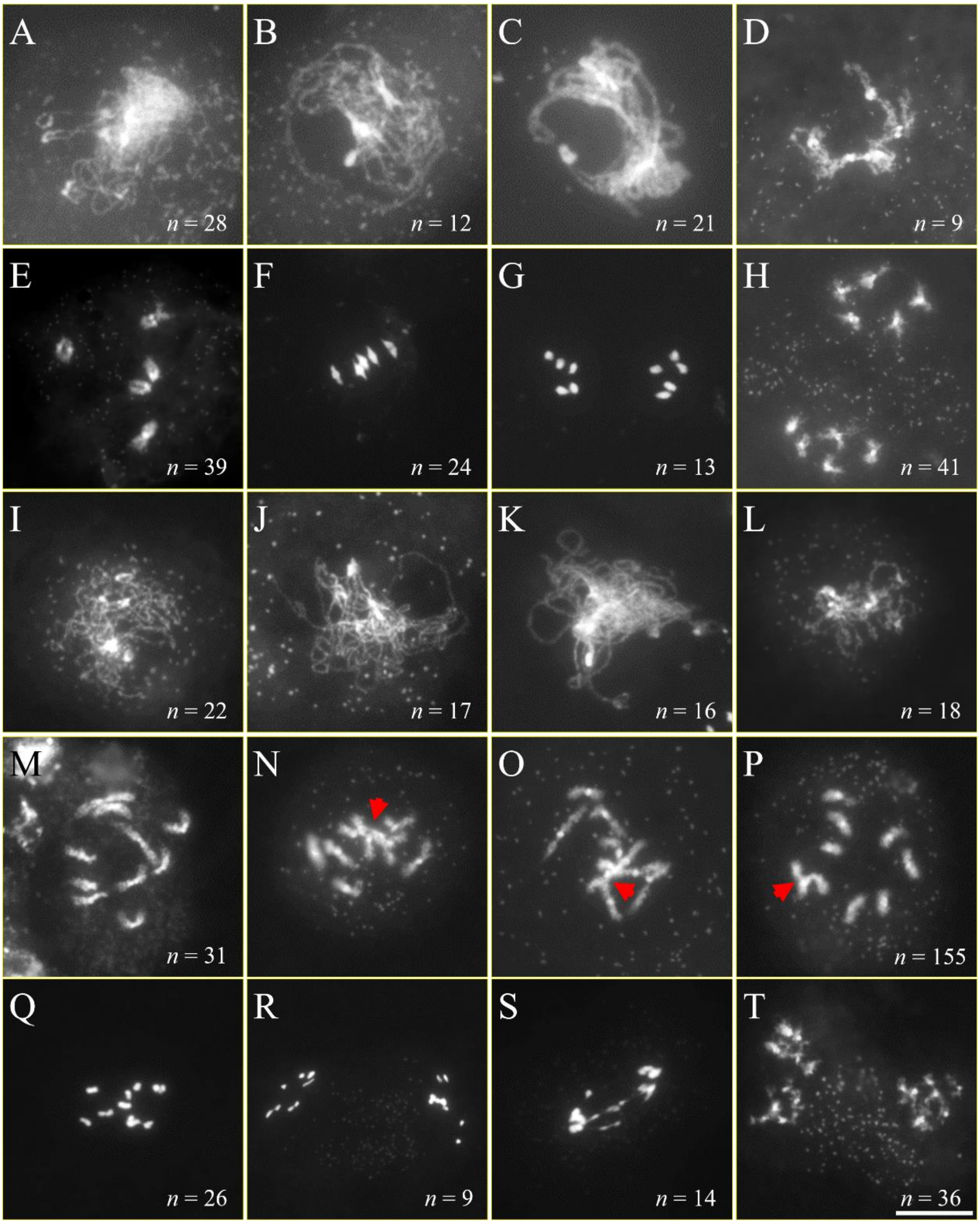


Supplemental Figure S1. DAPI-staining of meiotic chromosomes in heat-stressed wild-type Col-0 plants. A-H, Meiotic spread of Col-0 meiocytes at leptotene (A), zygotene (B), pachytene (C), diplotene (D), diakinesis (E), metaphase I (F), anaphase I (G) and interkinesis (H) stages under normal temperature conditions. I-T, Meiotic spread of heat-stressed Col-0 meiocytes at leptotene (I), zygotene (J), pachytene (K), diplotene (L), diakinesis (M-P), metaphase I (Q), anaphase I (R and S) and interkinesis (T) stages. Red arrows point to abnormal chromosome interactions. *n* indicates the number of cells with corresponding phenotypes. Scale bar = 10 µm.

*Supplemental figure S2.*

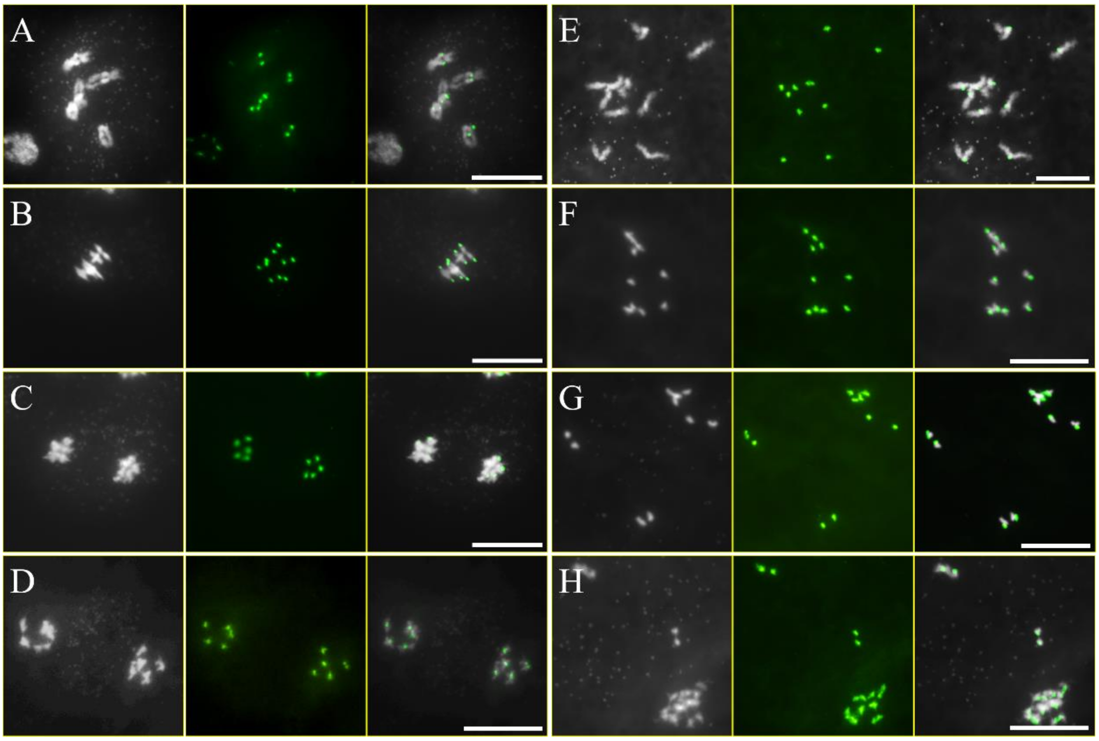


Supplemental Figure S2. FISH analysis of chromosome dynamics in heat-stressed wild-type Col-0 plants. A-D, Chromosomes at diakinesis (A), metaphase I (B), anaphase I (C) and interkinesis (D) stages in Col-0 plants under normal temperature conditions. E-H, Chromosomes at diakinesis (E), metaphase I (F), anaphase I (G) and interkinesis (H) stages in heat-stressed Col-0 plants. Chromosomes are stained by DAPI indicated by white, and centromeric 180 bp repetitive regions are labeled by a centromere-specific cen180_oligo2/5/6 probe shown by green. Scale bars = 10 µm.

*Supplemental figure S3.*

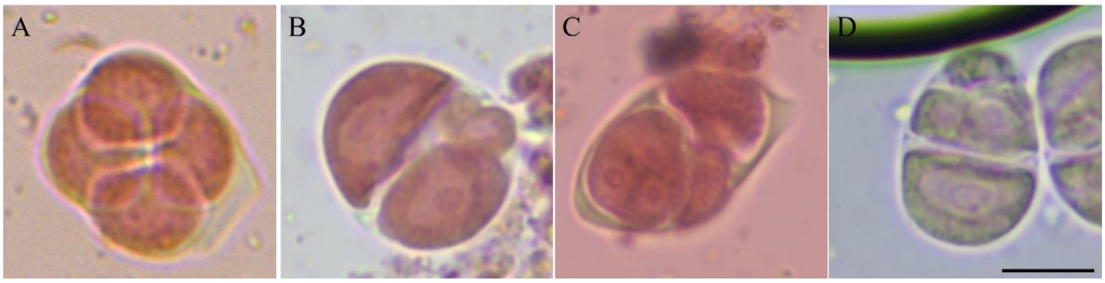


Supplemental Figure S3. Heat stress induces abnormal tetrad-staged PMCs. A-D, Meiotic products at tetrad stage produced by Col-0 plants incubated at normal temperature (A) and at high temperature conditions (B-D). Scale bar = 10 μm.

*Supplemental figure S4.*

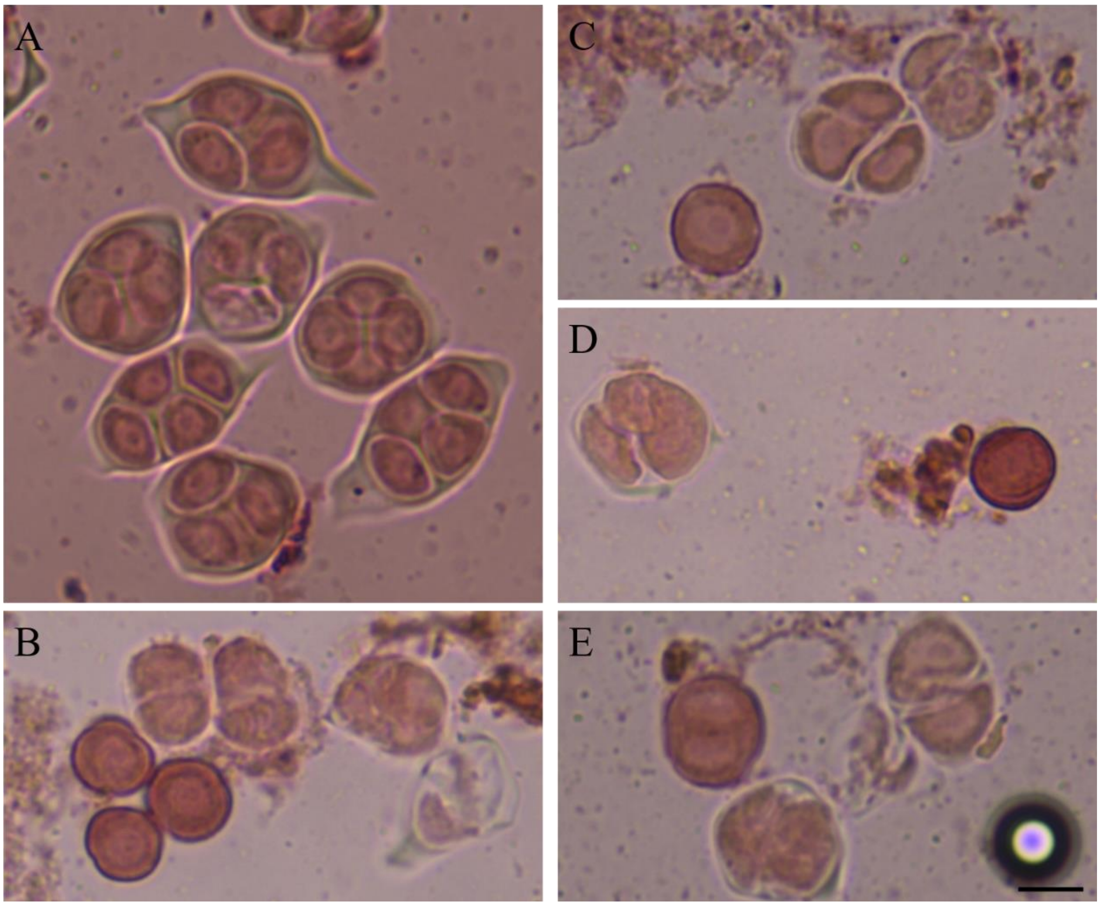


Supplemental Figure S4. Meiotic products of heat-stressed wild-type Col-0 plants stained by orcein. A-E, Meiotic products produced by Col-0 plants under control (A) and high temperatures (B-E). Scale bar = 10 μm.

*Supplemental figure S5.*

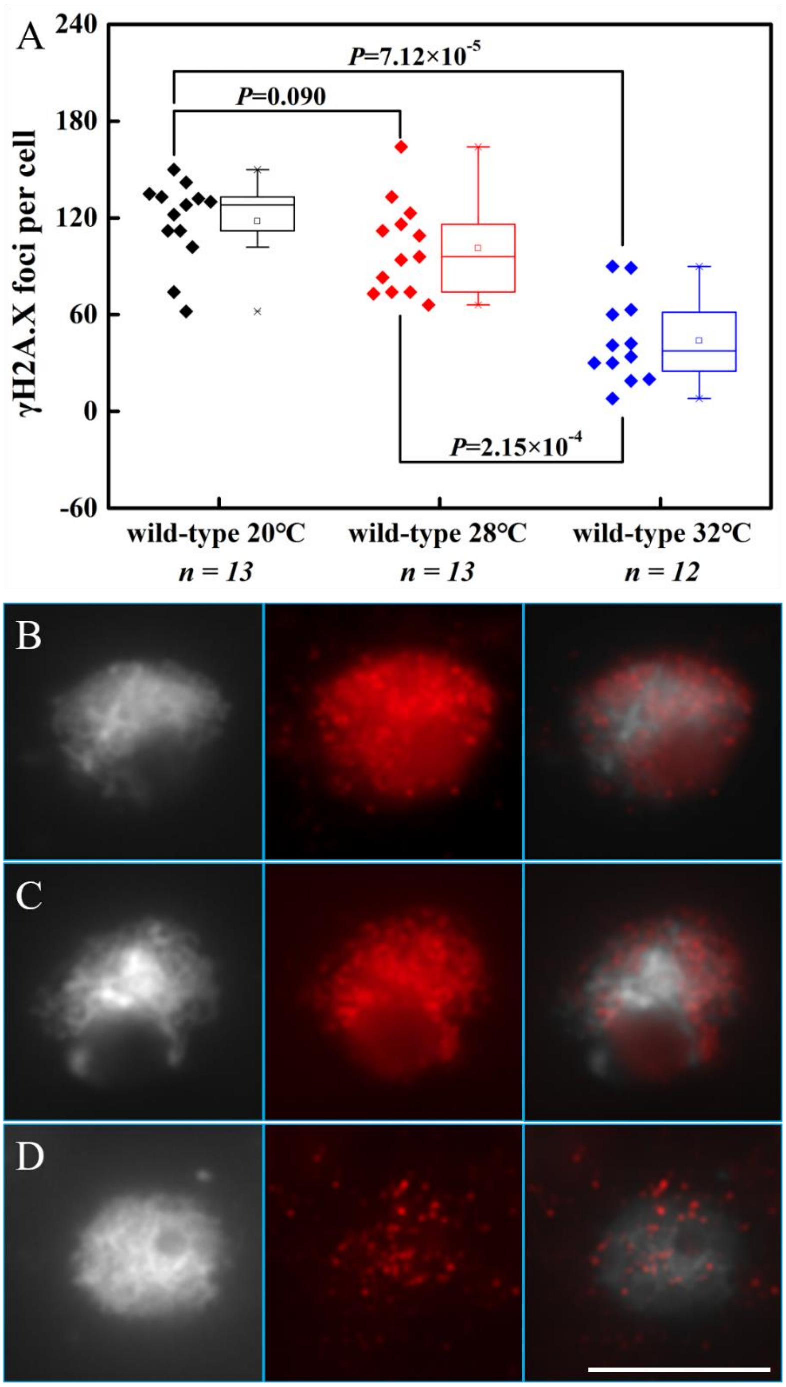


Supplemental Figure S5. Quantification of γH2A.X in wild-type Col-0 plants incubated at 20°C, 28°C and 32°C conditions. A, Graph showing the quantification of γH2A.X in wild-type Col-0 plants incubated at 20°C, 28°C and 32°C conditions. B-D, Localization of γH2A.X in wild-type Col-0 plants incubated at 20°C (B), 28°C (C) and 32°C (D) conditions. Two-side Mann-Whitney *U* test was performed. *n* indicates for the number of cells used for quantification. White: DAPI-stained chromosomes; Red: γH2A.X. Scale bar = 10 μm.

*Supplemental figure S6.*

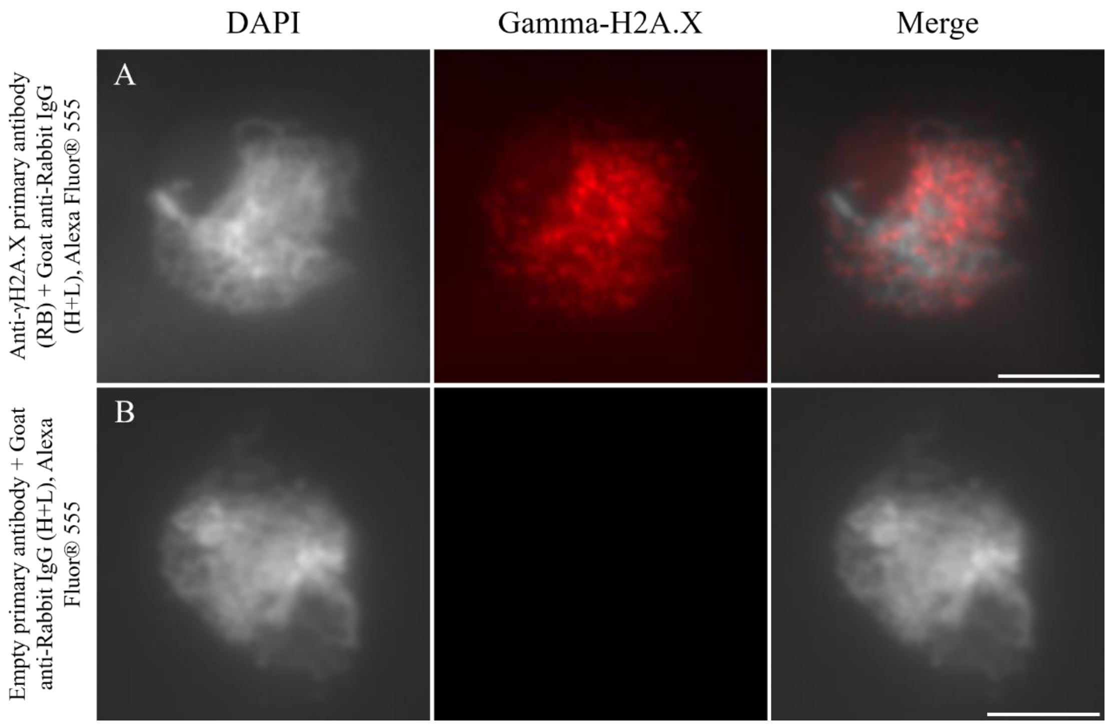


Supplemental Figure S6. Specificity test of anti-γH2A.X antibody. A, Immunolocalization of γH2A.X foci on wild-type zygotene chromosome. Rabbit anti-γH2A.X antibody was used and immunized by the secondary antibody Goat anti-Rabbit IgG (H+L), Alexa Fluor® 555. B, Immunolocalization of γH2A.X foci on wild-type zygotene chromosome. The secondary antibody Goat anti-Rabbit IgG (H+L), Alexa Fluor® 555 was directly added to samples. Scale bars = 10 μm.

*Supplemental figure S7.*

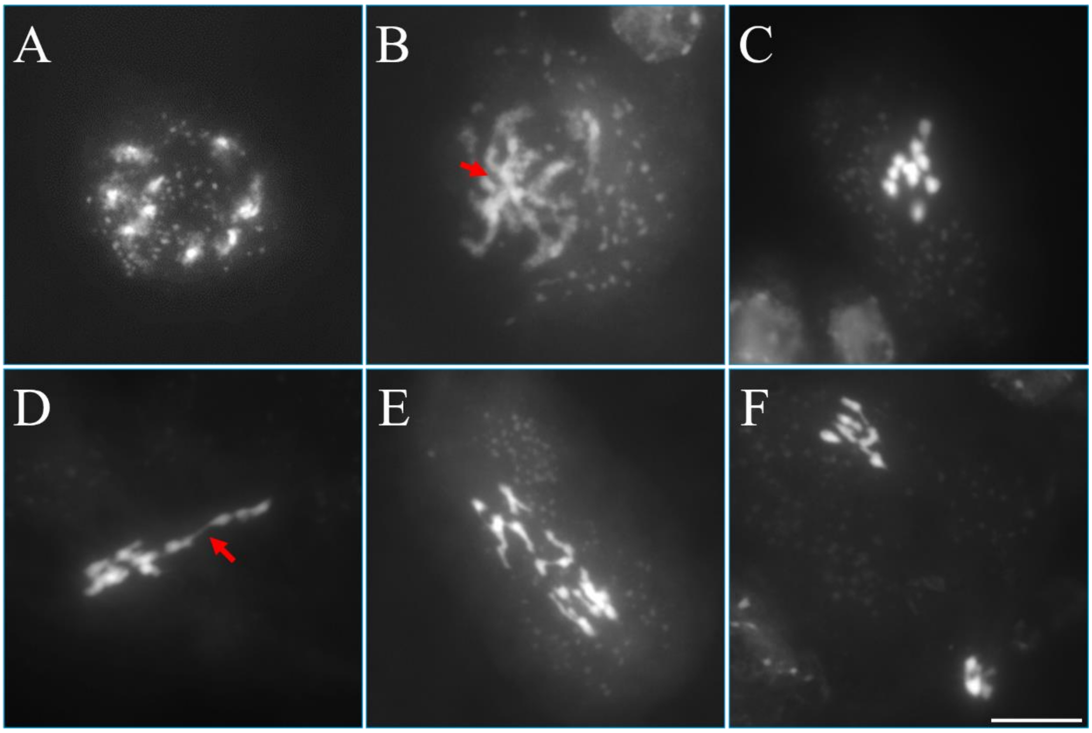


Supplemental Figure S7. Meiotic spread in the *spo11-1-1* mutant under control temperature. A-F, Diakinesis- (A and B), metaphase I- (C and D) and anaphase I-staged (E and F) PMCs in the *spo11-1-1* mutant under normal temperature conditions. Red arrows indicate irregular interactions between univalent. Scale bar = 10 μm.

*Supplemental figure S8.*

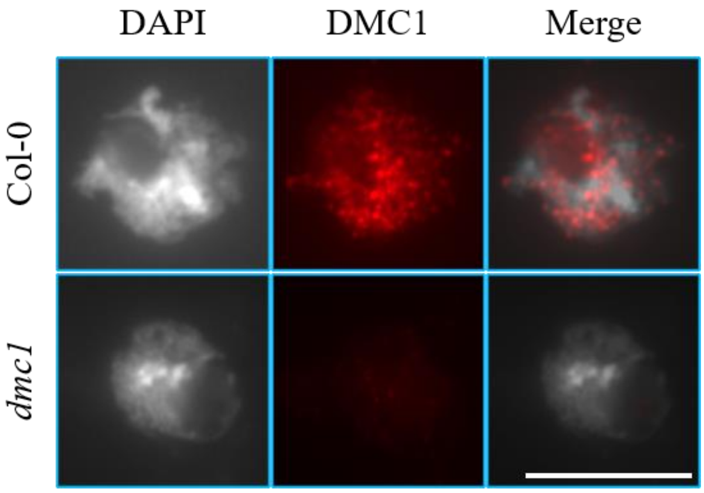


Supplemental Figure S8. Immunolocalization of DMC1 in the *dmc1* mutant. White: DAPI-stained chromosomes; Red: DMC1. Scale bar = 10 µm.

*Supplemental figure S9.*

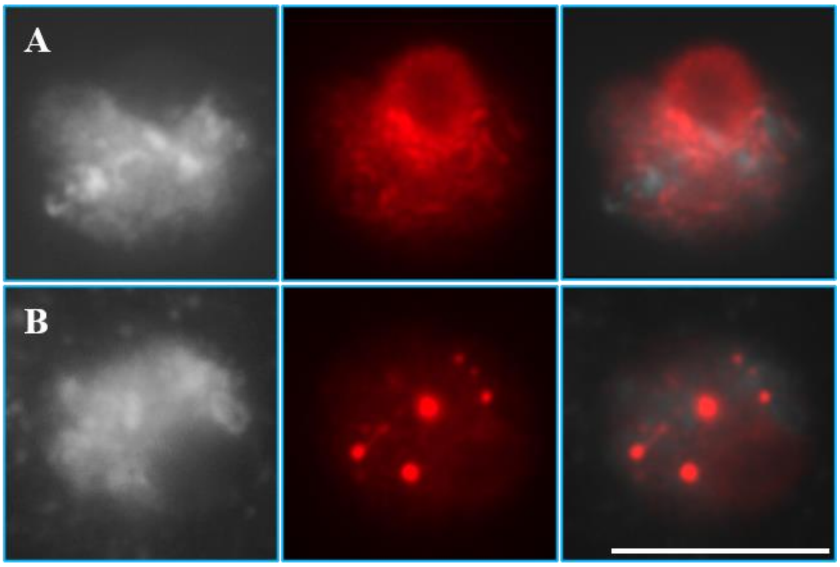


Supplemental Figure S9. Immunolocalization of ASY4 in the *syn1* mutant under normal temperature conditions. White: DAPI-stained chromosomes; Red: ASY4. Scale bar = 10 µm.

Supplemental Table S1. Primers used in this study.

| Primers | Sequences (5′→3′) | Purpose |
|---|---|---|
| *EF1αA4*-qRT-F | AGGCTGGTATCTCTAAGGATGGTCA | *EF1αA4*-specific primers for qRT-PCR |
| *EF1αA4*-qRT-R | GGATTTTGTCAGGGTTGTATCCG | |
| *HSFA1a*-qRT-F | AATGCGAACTCTTTACCGCCCCC | *HSFA1a*-specific primers for qRT-PCR |
| *HSFA1a*-qRT-R | AGCCATTGAACCTTGACCCTGAG | |
| *ASY1*-qRT-F | CCAGATTACGAGCCACCTTTCTT | *ASY1*-specific primers for qRT-PCR |
| *ASY1*-qRT-R | TGATCCAGTCCTTCACCCTTGCT | |
| *SYN1*-qRT-F | ATCCGTCGGTTCCGATGGCTCTT | *SYN1*-specific primers for qRT-PCR |
| *SYN1*-qRT-R | CTCGGCATCAGCTTGATGGAACT | |
| *ZYP1A*-qRT-F | GGGTTTCCGGCGATGAAGAGTTT | *ZYP1A*-specific primers for qRT-PCR |
| *ZYP1A*-qRT-R | ATCCTGAACTTGGGAAGCCAAAT | |
| *RAD51*-qRT-F | GTTGCCTATGCGAGGG | *RAD51*-specific primers for qRT-PCR |
| *RAD51*-qRT-R | ATCGGCTTAAATTGGGGA | |
| *SPO11-1*-qRT-F | CAGCTTTCAAAGCACAATCCATTGTG | *SPO11-1*-specific primers for qRT-PCR |
| *SPO11-1*-qRT-R | GATATCCTCTTCCTGTGATAACAATG | |
| *DMC1*-qRT-F | GGAGGGAATGGAAAAGTG | *DMC1*-specific primers for qRT-PCR |
| *DMC1*-qRT-R | GCAACGTTGAACTCCTCTGCAAT | |
| *ATM*-qRT-F | GATGGCCATGAGGCATTATT | *ATM*-specific primers for qRT-PCR |
| *ATM*-qRT-R | GCGGCAATGATTAATTCCCTCTGAC | |
| *ATR*-qRT-F | GTTGCTGCGTTCCAACTCAAGG | *ATR*-specific primers for qRT-PCR |
| *ATR*-qRT-R | TGCGGGAGGATGTTGAATGTAAGG | |
| *ASY3*-qRT-F | GATTTAGTTCTGTCTGATCCGTCG | *ASY3*-specific primers for qRT-PCR |
| *ASY3*-qRT-R | GTTTTGAAGAGCCATAGCGAAC | |
| *ASY4*-qRT-F | CTAAGAGAACTCGGCCAGAG | *ASY4*-specific primers for qRT-PCR |
| *ASY4*-qRT-R | TTTGACTTCACCTCAGTGGA | |
| *OSD1*-qRT-F | GTGGTGGTTTGTTGCCTTCTTGGT | *OSD1*-specific primers for qRT-PCR |
| *OSD1*-qRT-R | TCAACTTCTGGATCTCCGCCATCA | |
| *PRD1*-qRT-F | GACACAGAGAGATGATGTTCGCTTG | *PRD1*-specific primers for qRT-PCR |
| *PRD1*-qRT-R | GGTGTACTCTCTTGAGATATGTAGTGG | |
| *PRD2*-qRT-F | CCCGTTGTTATGTTTCTTCTG | *PRD2*-specific primers for qRT-PCR |
| *PRD2*-qRT-R | CAGACAGTTCCAGGAGAAAATC | |
| *PRD3*-qRT-F | GATCAGCAGAACCACAAGCATCTC | *PRD3*-specific primers for qRT-PCR |
| *PRD3*-qRT-R | CCGTTGCTCAAGCTCTTCACTGAT | |
| *TAM*-qRT-F | AATCCGATTCCTCGTCCGAACT | *TAM*-specific primers for qRT-PCR |
| *TAM*-qRT-R | GTCAGATTGAGTAGGAGAGGCAT | |
| *DMC1*-LP | GACTCATTGTTGCTTGATCCC | Genotyping the *dmc1* T-DNA mutant (SAIL_170_F08) |
| *DMC1*-RP | TCCACTCGGAATAAAGCAATG | |
